# Histopathological and virus-associated features of Boid Inclusion Body Disease in *Boa constrictor*

**DOI:** 10.64898/2026.11.03.764006

**Authors:** Maurice Bogner, Eva Dervas, Simon De Neck, Jussi Hepojoki, Francesca Baggio, Udo Hetzel, Anja Kipar

## Abstract

Boid inclusion body disease (BIBD) is a reptarenavirus-induced fatal disease of boid snakes that, while diagnosed with ease due to the pathognomonic intracytoplasmic inclusion bodies (IB), is not yet fully characterized pathomorphologically and with regards to its pathogenesis. The present study retrospectively investigated a random cohort of 50 captive male and female boa constrictors of variable age (neonate to 25 years) for histopathological changes and *in situ* viral protein (nucleoprotein, NP, and glycoprotein 2, GP2) expression patterns. IB formation and NP expression were scored semiquantitatively. The vast majority of snakes (42/50; 84%) exhibited histopathological changes; these mainly (39/42; 93%) represented inflammatory processes that most frequently affected the gastrointestinal tract. Overall, IB and NP scores were significantly higher in the more recent cases (2020-2025) compared to the older cases (2000-2003). NP scores did not differ between age groups or sex, but were lower in animals with clinical signs and higher in those with histopathological changes. No statistically significant association was found between NP scores and inflammatory processes, neither in general nor in individual organs. GP2 expression, as an indicator of virion assembly, was less consistent than NP expression and generally restricted to individual cells with and without IBs. There was no histological evidence of overt damage in infected cells. Jointly, the results indicate different phases of restrictive and systemic viral replication that are not directly associated with pathological processes and/or clinical signs.

## Introduction

Boid inclusion body disease (BIBD) is a transmissible disease of snakes from the families *Boidae* and *Pythonidae* and was first reported in captive collections in the late 1970s.^1^ BIBD has gained the name from its morphological hallmark, cytoplasmic eosinophilic inclusion bodies (IB) that accumulate in a broad variety of cell types including blood cells.^1–3^ In fact, the gold standard and minimal intervention *intra vitam* diagnostic approach is the examination of May-Grunwald stained blood smears for the presence of cytoplasmic IB within blood cells, and in particular erythrocytes.^4–6^

The most striking clinical indications reported in animals with BIBD are neurological signs,^6–10^ such as head tremor, opisthotonos (“stargazing”), torticollis and disorientation. Affected animals are also described as unable to right themselves when placed in dorsal recumbency, eventually progressing to flaccid paralysis.^1,5,11^ However, CNS signs seem to be rare with BIBD in recent years (personal observation) consistent with a recent report.^12^ In agreement with this, a substantial proportion of animals carrying IB within their blood cells and, per definition, affected by BIBD, do not show any overt clinical signs,^6–10^ and in a study where a large breeding collection of apparently healthy boas was screened for BIBD, the only clinical parameter found to be altered was the body weight in female snakes that was lower when the animals were affected by BIBD.^6^

A novel group of arenaviruses, later assigned to the genus *Reptarenavirus*,^13^ were identified as the causative agents of BIBD in the early 2010s.^2,3,14^ Reptarenaviruses are enveloped and have a bisegmented ambisense single-stranded RNA genome.^15^ The large (L) segment encodes the RNA-directed RNA polymerase (RdRp) and the Z protein, a matrix protein that in the case of mammarenaviruses regulates viral replication and facilitates virion assembly and budding.^15^ The small (S) segment encodes the nucleoprotein (NP) and glycoprotein precursor (GPC). Due to the ambisense coding strategy of arenaviruses, NP and RdRp mRNAs are transcribed directly from the genomic S and L segments, respectively, and can therefore be synthesized as the first proteins upon release to host cell cytoplasm. In contrast, GPC and Z protein mRNAs are transcribed from the corresponding antigenomic segments, requiring a prior round of viral RNA replication.^15^ NP is the main component of the IB in infected cells.^2–4^ GPC is co- and post-translationally cleaved into a signal peptide (SP), glycoprotein 1 (GP1) and 2 (GP2). In mammarenaviruses and hartmaniviruses, the SP is retained as a stable signal peptide (SSP) associated with GP1-GP2 heterodimers, whereas in reptarenaviruses it is likely not retained. Three glycoprotein heterodimers assemble to form the trimeric virion spike.^15–18^ GP1 mediates receptor binding, whereas GP2 drives the fusion between viral and endosomal membranes during virus entry.^15^

Observations from the work-up of the boa constrictors with BIBD that we received for diagnostic post mortem examinations suggest that, despite the wide distribution of the intracytoplasmic IB, affected animals exhibit a rather limited range of histopathological changes. Based on our recent findings, the intestines appear to be most frequently affected, as a heterophilic colitis without histological evidence of an additional intestinal infectious etiology was the most common lesion in snakes with BIBD.^12^ Considering that the clinical presentation of BIBD may have become more subtle, less specific, and less frequently characterized by neurological signs than originally described,^1^ we hypothesized that the pathological manifestation of BIBD may also have evolved. We also hypothesized that clinically apparent BIBD represents a more advanced stage of disease, with increased lesion severity, broader tissue tropism of infection, greater IB burden, and enhanced viral antigen expression compared with clinically healthy BIBD-positive snakes, and wondered whether these pathological features vary according to the age and sex of affected animals. To address these hypotheses and to reveal potential differences in parameters over time, a large randomly selected cohort of captive boa constrictors with confirmed BIBD, collected over a period of just over 25 years, and with a wide range of clinical presentations, different age and sex, was retrospectively subjected to an in-depth histological analysis.

Earlier *in vitro* studies of our group have investigated the viral persistence in permissive boa constrictor cells and detected consistent NP expression but a significant decline in GP2 expression during persistent infection.^19^ This led us to hypothesize that IB formation would be prevented by constant egress of NP when GPs are expressed and virions formed, but when GP expression declines during persistent infection the NP accumulation would generate IB. To address this hypothesis, we examined the snake cohort in the present study for the distribution of viral NP (as indicator of replication) and GP2 (as indicator of virion formation), and correlated the findings with all other parameters and clinical outcomes.

## Material and methods

### Study animals

The study was undertaken on 50 captive *Boa constrictor* with confirmed BIBD that had died with (animals no. 3 and 24) or were euthanized due to the disease (all other animals) and were subjected to a full post mortem examination for diagnostic purposes between 2000 and 2025. Animals were either individually owned snakes from private households (n=19), originated from different breeding collections in Germany or Switzerland (n=23), or were from a reptile shelter in Switzerland that takes in reptiles from private owners or breeders and attempts to rehome them (n=8; cases 37, 40, 41.1-41.5, 42). Election for euthanasia was either due to severe clinical disease or because a diagnosis of BIBD had been made (based on the presence of the characteristic intracytoplasmic IBs in cells in a peripheral blood smear). As the owners wanted to clear their colony/shelter from BIBD, they submitted the snakes for euthanasia and post mortem examination, to confirm BIBD and to determine any other, potentially relevant concomitant pathological conditions.

All snakes submitted alive for euthanasia and subsequent diagnostic post mortem examination were euthanized by veterinarians according to the ASPA, Animals (Scientific Procedures) Act 1986, schedule 1 (appropriate methods of humane killing, http://www.legislation.gov.uk/ukpga/1986/14/schedule/1) procedure. The post mortem examination was performed upon the owners’ request. The use of the diagnostic samples for research purposes was approved by the institutional review board (MeF-Ethik-2024-01). Also, the terms of service to which owners agree when submitting an animal for a diagnostic post mortem examination include the permission to make use of material from the examination for both teaching and research.

### Histological and immunohistochemical examinations

Immediately after necropsy, organ samples were collected for diagnostic histological examination. These were fixed in 10% neutral buffered formalin for 1-3 days, then trimmed and routinely embedded in paraffin wax. Consecutive sections (2-3 µm) were prepared and stained with hematoxylin and eosin (HE) for histological examination or subjected to immunohistochemistry (IHC) for the detection of reptarenavirus nucleoprotein (NP) or glycoprotein (GP2).

In all animals, HE stained sections served to confirm BIBD based on the presence of the typical IBs in cells in the different tissues, and to determine any histopathological changes. The following main organs were available for consistent histological examination: lung (n=45), stomach (n=41), small and large intestine (n=50), liver (n=49), pancreas (n=43), kidney (n=49), spleen (n=41), central nervous system (n=46; brain: 44, spinal cord: 23).

The abundance of IBs was assessed in HE stained sections using a semi-quantitative grading system applied to the tissues/organs that could be assessed most reliably, i.e. brain and/or spinal cord, pancreas and spleen. Scoring was performed by an investigator blinded to age, sex and origin of the animals. Slides were evaluated sequentially, with IB scores assigned separately for the brain, pancreas, and spleen based on the histological assessment of IB abundance. To promote consistency in scoring, previously examined slides of the corresponding organ were consulted as visual references whenever needed. Values ranged from 0 (no evidence of IBs), over 0.5 (few IBs) to 3 (very numerous IBs), in 0.25 steps.

For IHC, the horseradish peroxidase method was applied. Sections were routinely deparaffinized. Heat induced antigen retrieval was performed by incubating the slides in citrate buffer (pH 6.0; Agilent Dako) for 20 min at 98 °C in a pressure cooker. Sections were then incubated overnight at 4 °C with the respective primary antibody at a dilution of 1:6,000 in Antibody Diluent (Agilent Dako). Previously described rabbit antisera raised against *Escherichia coli* produced multiepitope recombinant proteins designed for reptarenavirus nucleoprotein (NP) and glycoprotein 2 (GP2) were utilized as primary antibodies to detect reptarenavirus antigens.^19,20^ This followed by blocking of the endogenous peroxidase activity with Peroxidase Blocking Solution (Agilent Dako) for 10 min at room temperature. Antibody binding was detected by incubating the slides with the EnVision+System-HRP anti-rabbit polymer (Agilent Dako), followed by visualization with DAB+ Substrate Buffer (Agilent Dako), each for 30 min at room temperature. These incubations were carried out in an autostainer (Agilent Dako). Slides were counterstained with hematoxylin (Gill2, Carl Roth). Formalin-fixed and paraffin-embedded tissue samples from snakes with confirmed BIBD (positive control) and from reptarenavirus (RT-PCR) negative snakes (negative control) served as general controls, and sections subjected to the above protocols where the primary antibodies were omitted served as case-specific negative controls.

IHC was performed on the majority of inner organs, with the following tissues being most consistently available for IHC: brain (n=44), spinal cord (n=20), liver (NP: n=45; GP2: n=41), pancreas (NP: n=42; GP2: n=39), kidney (NP: n=41; GP2: n=29) and spleen (n=39).

Sections stained by IHC for viral NP expression, representative of IBs, were investigated to determine the distribution of NP, i.e. the tissues and cells that harbored IBs, and served to semi-quantitatively assess the extent of viral NP expression. The latter was restricted to sections from brain and/or spinal cord, pancreas and spleen, since the presence of endogenous pigments (i.e. bile and hemosiderin) impaired attempts at quantification of DAB deposition (and hence viral NP expression) in sections from liver and kidney in particular. A scale from 0 (absent) to 3 (strongly positive) was applied, with intermediate values (0.25 steps). For those individuals where both spinal cord and brain tissue sections were available, a combined semi-quantitative value was determined for the CNS, based on the arithmetic mean. If either brain or spinal cord was available, the CNS value was based on this.

Since the staining for GP2, as representative of virion production, was not consistent and generally weak and limited to individual cells, scoring was not attempted; instead, individual staining results were recorded (Supplemental Table S2).

### Statistical analyses

Statistical analysis was performed using R (v4.5.2)^21^ with RStudio (v2026.01.0 Build 392, Posit, Boston, USA). This based on the semiquantitative scores applied to sections of CNS, pancreas and spleen stained for viral NP and stained with HE (IBs). For comparative analyses considering the age, animals were allocated to age groups, summarized as perinatal (≤2 months; n=5), subadult (3 months <3 years; n=21), or adult (≥3 years; n=24). The cutoff age of 3 years was used to distinguish subadult from adult boas, consistent with our previous studies.^22,23^ This threshold approximately corresponds to the onset of sexual maturity and reproduction, which have been shown to be associated with significant changes in hematological and biochemical parameters, as well as histological alterations in hemolymphatic tissues. The following comparisons were conducted: 1) older material (cases from 2000-2003; n=21) and recent material (cases from 2020-2025; n=20): for both NP and HE scores; 2) age groups; 3) sex (female, male); 4) clinical signs reported or not; 5) presence/absence of histological changes; 6) presence/absence of inflammatory processes; 7) presence/absence of gastritis/enteritis/colitis, pneumonia/tracheitis, hepatitis or nephritis; 8) presence/absence of splenitis, macrophage rearrangement and granular cell infiltration in relation to the overall NP score and the splenic NP score; 9) presence/absence of inflammatory processes in more than one organ (tracheitis/pneumonia and osteomyelitis/myositis were considered as one entity, respectively).

## Results

### Study population and clinical signs

The study included 50 captive boa constrictors with confirmed BIBD, sampled between 2000 and 2025, with a broad age range (perinatal to 25 years). Of these, 5 were categorized as perinatal/juvenile (up to 2 months), 21 as subadult (<3 years), the remaining 24 as adults (≥3 years); 15 were male and 19 female. For 16 animals, the sex was not known (not recorded or not identifiable due to the lack of gonadal differentiation associated with young age).

Two animals in the study (cases 3 and 24) had died spontaneously; 16 had been euthanised due to clinical signs, 9 of these after the BIBD diagnosis. Twenty-four animals were euthanized after BIBD diagnosis based on a blood smear (presence of IB in blood cells), without clinical signs. Five 4-month-old siblings (cases 41.1 to 41.5) as well as their mother (case 40) were euthanized after the mother had received the BIBD diagnosis. One neonate was euthanized due to a malformation (case 15; coelomic herniation), and two adults after several unsuccessful rehoming attempts (cases 40 and 43). Most snakes with clinical signs (n=15, including one animal found dead) had been examined in the years 2000-2003. Clinically, poor body condition/emaciation was most frequently reported (13 of the 16 animals from which clinical details were provided; 81.25%), 4 of these animals had also shown weakness or apathy. In five animals (cases 1.3, 4, 5, 10 and 22.1; 31.25%), clinical signs indicative of CNS disease were noticed; these included headshaking, anisocoria and a lack of righting reflex but were otherwise not specified. In 5 animals, examined in 2000 and 2001 (cases 2, 4, 5, 7 and 10; 31.25%), the alimentary tract was clinically affected; fibrinonecrotic stomatitis (n=2), regurgitation and emesis, diarrhea, and coprostasis (each n=1) were reported. Three animals (cases 6, 10, 24; 18.75%) exhibited respiratory signs; in two of these, rhinitis was reported. Another two animals (cases 7 and 13; 12.5%) were diagnosed with a suppurative myositis and osteomyelitis, in one animal together with an abscess in the Jacobson’s (vomeronasal) organ. Individual animal data are provided in Supplemental Table S1.

### Histopathological changes

The histological examination revealed frequent, though limited pathological changes. These mainly represented inflammatory or degenerative processes. The <u>gastrointestinal tract</u> was most frequently affected (32/50; 64%) and exhibited exclusively inflammatory changes, mainly affecting the small and/or large intestine (n=31). The inflammatory infiltration was mainly mixed cellular, in the majority with heterophil dominance (n=18; Supplemental Figure S1a). In two and one cases, respectively, a fibrinonecrotic (cases 7, 24) and ulcerative (case 4) inflammation was observed. The presence of intralesional bacteria indicated a bacterial etiology; however, as no bacteriological examination was performed, comments on the causative organisms cannot be made. In each two cases, a (pyo)granulomatous (cases 10, 12) and an almost purely lymphocytic (cases 1.1, 8) infiltrate was seen. The inflammatory infiltrates were generally predominanly located in the submucosa (Supplemental Figure S1a). Six animals (cases 2, 4, 8, 12, 25, 40; 6/39, 15.4%) displayed lymphocyte aggregates in the gastric mucosa and/or submucosa (Supplemental Fig. S1b), interpreted as chronic lymphocyte dominated gastritis, in three cases alongside inflammatory processes in the intestines.

The <u>spleens</u> also frequently exhibited histological changes (19/41; 46.34%). These were mainly represented by a rearrangement of macrophages, a known splenic feature in boas with BIBD,^12^ which was seen in 14 cases, almost consistently (12/14) together with inflammatory processes in other organs or the spleen, and in 7 animals alongside granular cell infiltrates in the spleen (Supplemental Figure S1c), another feature recently reported in spleens of boas with BIBD.^12^ Splenitis was observed in 7 cases, in all but one animal (case 34) alongside inflammatory processes in at least one other organ. In most cases (5/7), the splenitis was of granulomatous nature, in one of these (case 24) with a clear necrotic component. The diagnosis of granulomatous splenitis was made when, different from the described macrophage rearrangement, well-demarcated nodular aggregates of macrophages, frequently with epithelioid appearance and occasionally surrounded and infiltrated by heterophils, were present (Supplemental Figure S1d). In the remaining two cases (cases 34 and 36) a heterophil-dominated splenitis was observed, in case 34 as the only inflammatory process.

In thirteen cases (13/45; 28.89%), the <u>lungs</u> were found to exhibit inflammatory processes. These were of overt chronic nature in 10 cases: In 7 animals, a lymphocyte-dominated interstitial pneumonia was seen (Supplemental Figure S1e), accompanied by chronic tracheitis in one case (case 32) and harboring intralesional nematodes (larvae and adults) in another (case 4). In another three cases, the lymphocytes were accompanied by heterophils which dominated the infiltrate in an additional case. In two animals, a granulomatous infiltrate was observed which in one case (case 5) was present alongside a fibrinonecrotic stomatitis and granulomatous hepatitis, likely representing a systemic bacterial infection; however, a bacteriological examination had not been performed.

In 12 snakes (12/49; 24.49%), inflammatory processes were observed in the <u>liver</u>. These were either of granulomatous nature (n=6; Supplemental Figure S1f, g; each one of these pyo- and lipogranulomatous), necrotic (n=2; cases 24 and 43), heterophil-dominated (n=2; cases 1.3 and 12), and in one animal (case 32) lymphocytic. In most cases, the hepatitis was observed alongside enteritis/colitis, indicating dissemination of intestinal bacteria to the liver. In two cases, it was accompanied by pneumonia, suggesting a systemic (bacterial) infection. In one case (case 11), multiple abscesses were detected; these would be consistent with bacterial spread from the intestines, however, they were the only histopathological lesion in this case. Three snakes exhibited hepatic lipidosis (cases 2, 21 and 37), in one case alongside poor body condition and clinical signs of regurgitation; another animal exhibited bile stasis in association with severe colitis (case 31), potentially reflecting reduced food intake associated with the underlying disease process.

In six snakes (6/49; 12.24%), all adult, the <u>kidneys</u> exhibited inflammatory changes. These were in the majority of chronic nature (5/6), represented by lymphocyte-dominated interstitial nephritis and pyelonephritis, partly with or dominated by fibrosis (Supplemental Figure S1h). In one animal (case 33), an acute heterophil-dominated nephritis with focal dystrophic mineralization was seen.

In four cases (cases 16, 37, 40, 41.2), the <u>thymus</u> showed histological changes. These were represented by epithelial cysts and occurred in animals of all ages (4 months, juvenile, 4 and 25 years), a finding in agreement with previous observations in boas.^12^ Further changes were a severe granulomatous salpingitis in an adult female (case 12), and nodular hyperplasia of the exocrine pancreas in an 11-year-old female (case 33).

Altogether, inflammatory processes were detected in a total of 39 animals (78%); in 21 of these (53.8%), they were present in more than one organ. Two organs were found to be affected in 14 animals, four animals showed inflammatory processes in three organs, and three in four organs. Different organ combinations were affected, in most (17/21) involving the intestine. In 6 animals, affected organs harbored intralesional bacteria (data not shown) which were not further examined. Detailed information on the histological changes is provided in Supplemental Table S1.

### Presence and distribution of cytoplasmic IB and in situ expression of reptarenavirus NP: potential relation to animal and disease parameters

In all animals, the typical cytoplasmic IBs were detected in some or all examined tissues.^1,3,5^ In HE stained sections, these were most consistently observed in brain and spinal cord (neurons, ependymal cells), kidney (primarily in tubular epithelial cells), liver (hepatocytes), pancreas (epithelial cells) and spleen (leukocytes) (Figure 1; Supplemental Figure S1h, S2); when detected in the gastrointestinal tract and lung, they were mainly visible in epithelial cells (Supplemental Figure S2). The IBs varied in size and shape, ranging from small dots in the cytoplasm to large, well-delineated round, ovoid or occasionally odd-shaped eosinophilic to amphophilic structures (Figure 1). There was no obvious pattern in size and number of IBs across organs/tissues or individual animals which was confirmed by the semi-quantitative scoring applied to CNS (brain and/or spinal cord), pancreas and spleen (Supplemental Table S2). However, the largest and unusually shaped IBs were detected in neurons in brain, spinal cord or spinal ganglia (Figure 1a).

**Figure 1.**
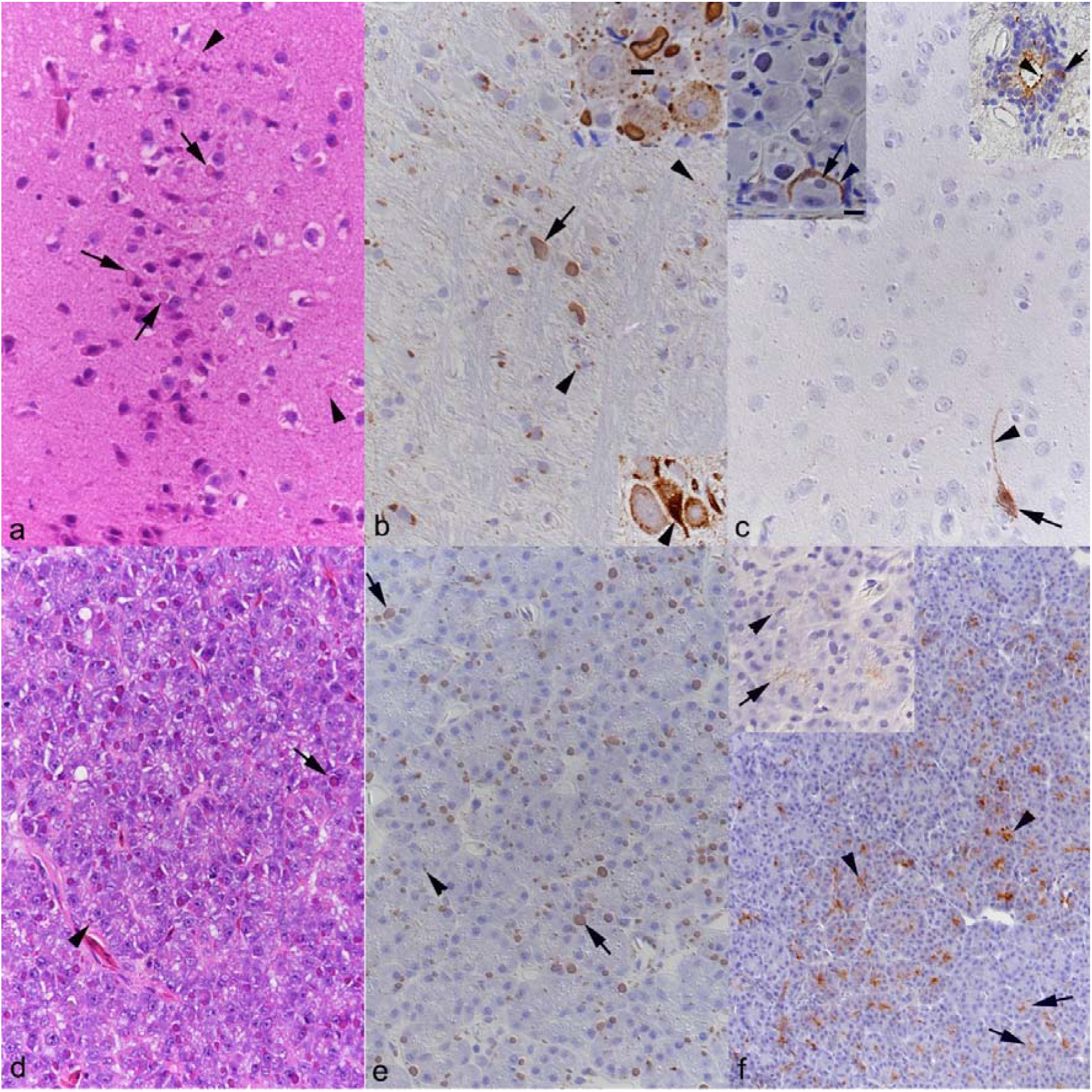
Inclusion body (IB) morphology and viral antigen expression, examplified in central nervous system (a-c) and pancreas (d-f). **a)** Brain, case 25 (male, 7 years). In the HE stained section, abundant variably sized delineated amphophilic IBs are seen in the cytoplasm of neurons (arrows). Several small IBs appear to be free in the neuroparenchym (arrowheads), likely representing IBs in neuronal processes. **b)** Brain, case 40 (female, 4 years). Viral NP expression is represented by well delineated, variably sized dots consistent with IBs seen in HE stained sections. Individual neurons exhibit one large IB with weakly stained center and more intensely labeled outer rim (arrow). Arrowheads indicate small IBs in neuronal cytoplasm or free in the parenchyma. Bottom inset: brain, case 43 (female, 12 years). Neuron with diffuse granular cytoplasmic NP expression (arrowhead). Top inset: spinal ganglion, case 41.5 (male, 4 months). Neurons with variably sized IBs, including one very large IB (bar = 10 µm) with weakly stained centre. Immunohistochemistry (IHC). **c)** Brain, case 43 (female, 12 years). Viral GP2 expression in individual neuron, with finely granular reaction in the entire cytoplasm (arrow) stretching into the neuronal process (arrowhead). Left inset: spinal ganglion, case 41.2 (male, 4 months). Reaction restricted to the cell periphery (arrowhead) in a neuron with IB (arrow). Right inset: spinal cord, case 41.4 (female, 4 months). GP2 expression in the cytoplasm (arrow) and along the luminal surface (arrowhead) of ependymal cells. IHC. **d, e)** Pancreas, case 40 (female, 4 years). Variably sized amphophilic cytoplasmic IBs in acinar epithelial cells in HE stained section (**d**), highlighted in sections stained for viral NP (**e**). Arrowheads: small IBs, arrows: large IBs. **f)** Pancreas, case 14 (neonate) and case 40 (inset). Viral GP2 expression is seen at the luminal surface (arrows) and as a cytoplasic reaction (arrowheads) in acinar epithelial cells, to a highly variable extent (inset: overall weak and scattered reaction). IHC.

In order to obtain a better picture of the extent and pattern of IB formation in the different tissues and animals, IHC for reptarenavirus NP, the main component of the IBs, was performed. This yielded cytoplasmic reactions consistent with IBs, with a staining pattern that generally mirrored the distribution of IBs seen in the HE-stained sections (Figure 1a, b). The NP staining reaction provided a sharper outline of the IBs, allowing determination of their size; this showed that the largest IBs had a diameter of more than 10 µm (Figure 1b). Interestingly, the larger IBs often exhibited a weak NP reaction, particularly in the center, often surrounded by a rim of more intense staining (Figure 1b). In some brains and/or spinal cords, individual or groups of neurons were found to exhibit a strong, diffusely granular cytoplasmic reaction that was not associated with defined IBs (Figure 1b).

The semiquantitative scoring of the overall NP staining intensity in CNS, pancreas and spleen did not suggest an organ or tissue specific staining pattern across the animals or differences in IB abundance in association with age or sex of the animals (Supplemental Table S2). However, we subsequently undertook detailed statistical analyses to test these observations.

Since we undertook the study on case material collected over the past 26 years, we investigated it for changes over time. In a first step, we examined it for potential effects of long-term paraffin storage on antigen preservation. For this purpose, we compared the semiquantitative NP scores obtained for the older archival cases (2000-2003; n=22) with those obtained for the more recent cases (2020-2025; n=22). Indeed, the staining scores were overall significantly higher in the recent material (NP: p<0.0001, Figure 2a). When the individual organs were compared, the difference was significant for CNS and spleen but not the pancreas (Supplemental Figure S3A). These results could indicate a partial loss of antigenicity in the older material (>20 years in the archive). However, since we could also score the IB quantity in the HE stained sections, the detection of which should not be affected by long term storage of paraffin blocks, we compared the two case groups for differences in the extent of IB formation. Surprisingly, this showed significantly higher overall scores in the more recent material (p<0.0001; Figure 2b), indicating both more extensive viral IB formation and NP expression in the more recent material, discarding a potential loss of antigenicity.

**Figure 2.**
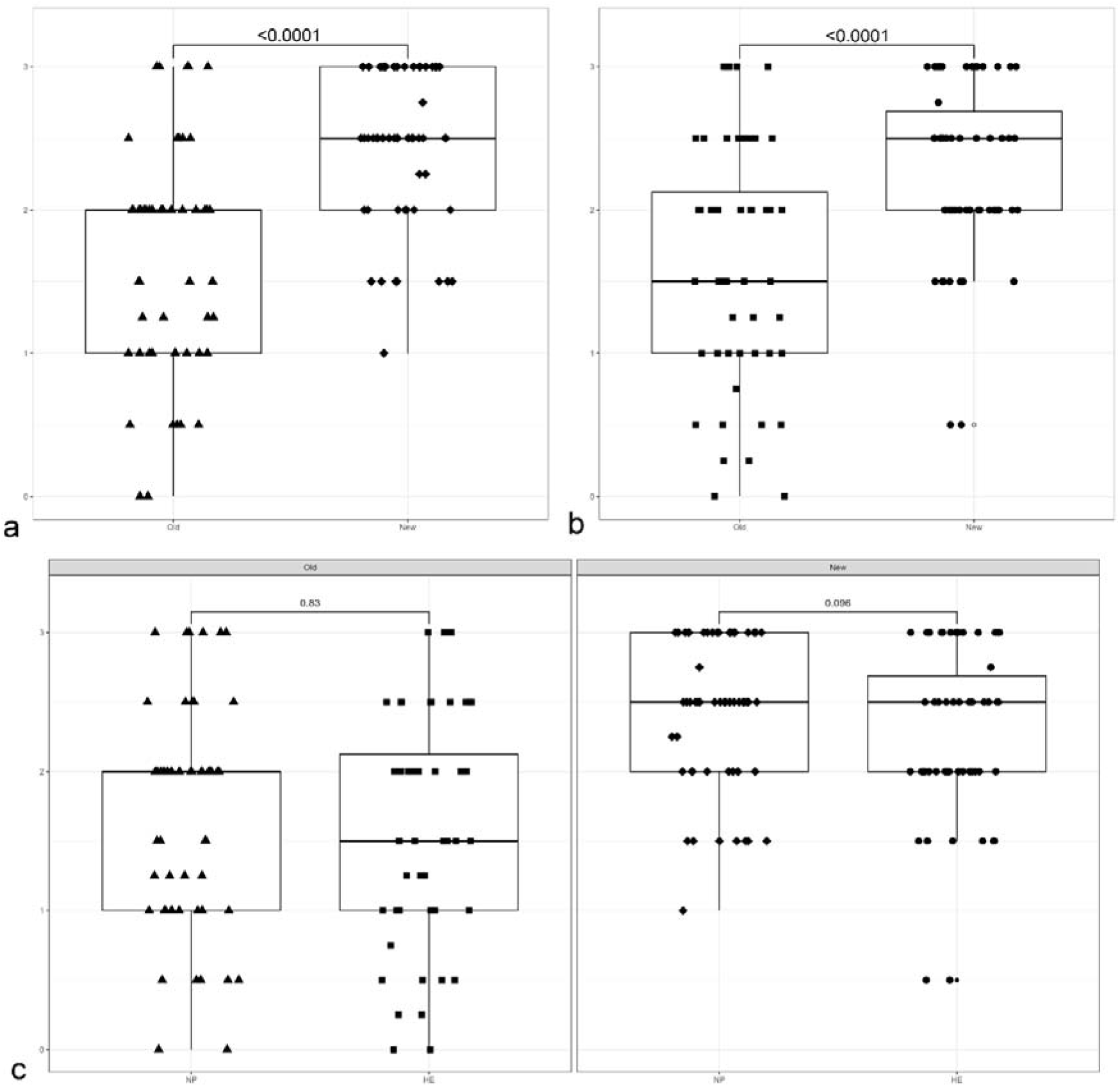
Comparison of mean overall NP expression scores (determined in IHC stained sections) and IB scores (determined in HE stained sections) in older archival material (2000-2003; n=22; “Old”) and more recent material (2020-2025; n=22; “New”), showing significantly higher overall NP scores (**a**) and IB scores (**b**) in the recent material. The NP and IB scores (**c**) did not differ significantly from each other in the older archival and the more recent material.

When the IB and NP scores of individual organs were compared, the difference between the older and more recent material was significant only in CNS and spleen, (Supplemental Figure S3). This could indicate a potential evolution of the disease, potentially also a virus-host adaptation, over time. We then compared the overall scores obtained from HE stained sections and from sections stained for NP for the two case groups; this did not reveal any significant differences in the NP vs the HE stain based scores within either group (Figure 2c). However, subsequent comparison of the scores in the three individual organs revealed significantly higher scores for NP than for IBs (HE stain based) in the CNS in the recent cases (Supplemental Figure S4). This could also indicate viral antigen expression without IB formation in neurons for which we found occasional IHC evidence, as shown in Figure 1b.

We subsequently undertook comparative assessments on all animals from the entire study cohort (2000-2025; n=50), using the NP expression scores as they reflect virus infection of the cell. First, **potential age-related differences in the extent of viral NP expression** were investigated. For this, the three age groups, perinatal/juvenile (≤3 months; n=5), subadult (3 months <3 years; n=21), or adult (≥3 years; n=24) were compared. Scores varied substantially within each group, without statistically significant differences between them (p=0.081; Figure 3a). Next, we investigated any **potential sex-related differences in viral NP expression**. For this, the data from animals with known sex were included (19 females; 15 males). Again, scores varied widely in both groups, and there was no significant difference in the mean scores between males and females (p=0.22; Figure 3b).

**Figure 3.**
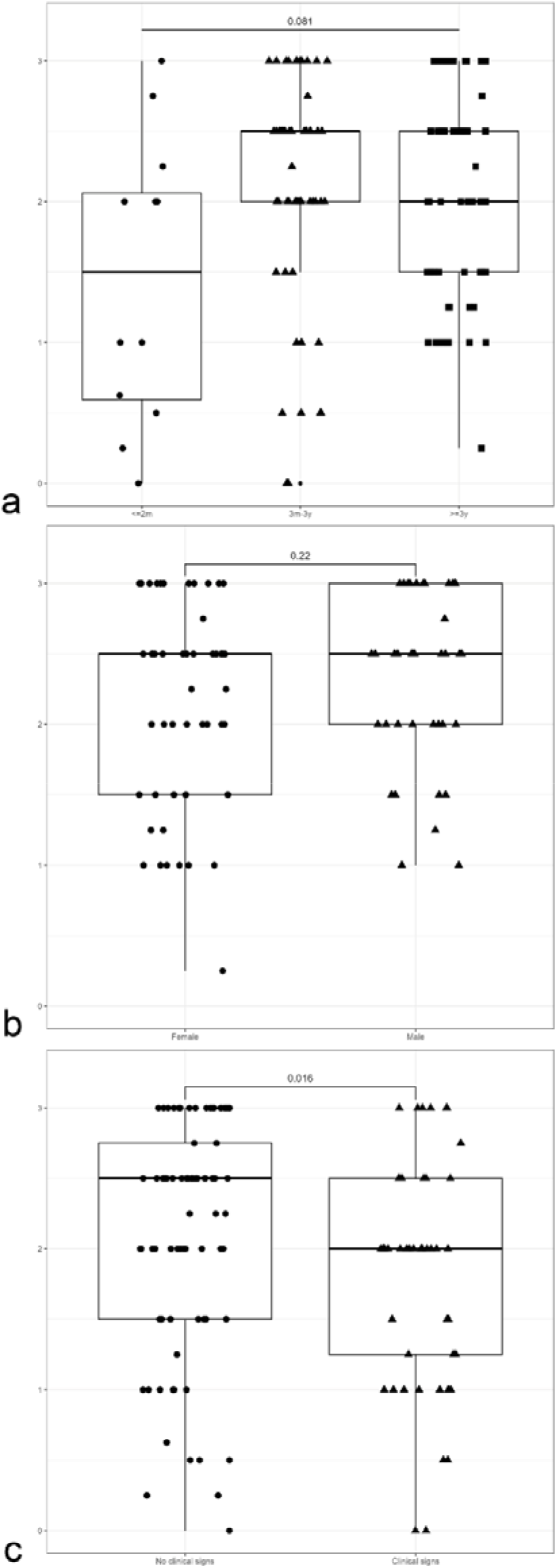
Comparison of mean overall NP expression scores (determined in IHC stained sections) does not show significant differences between (**a**) the three age groups, perinatal/juvenile (≤3 months; n=5), subadult (3 months <3 years; n=21), and adult (≥3 years; n=24); and (**b**) females (n=19) and males (n=15). Comparing animals without clinical signs (n=32) and with reported clinical signs or spontaneous death (n=18) (**c**) revealed significantly lower overall NP scores in the latter group. No: not observed; Yes: observed.

We then put emphasis on the **potential association of viral NP expression with disease processes**. In a first step, the **potential association with clinical signs** (including also spontaneous death) was investigated. Interestingly, this revealed significantly lower overall NP expression scores in animals with clinical signs (n=18; p=0.016) (Figure 3c). Considering that the majority of these animals (n=15) were in the older archival material, this may mirror the above described finding of an overall less extense NP expression in the older case material.

Since the majority of snakes had a range of histopathological changes, we next investigated the material for a **potential association between viral protein expression and the presence of histopathological changes** (detected in 42/50 animals). Indeed, while the scores varied substantially in both groups, mean scores were significantly higher in animals with histological lesions (p=0.016; Figure 4a). However, investigation of the **potential association with inflammatory processes**, as the dominant type of pathological processes, did not yield the same result for animals with inflammatory processes (p=0.051; Figure 4b).

**Figure 4.**
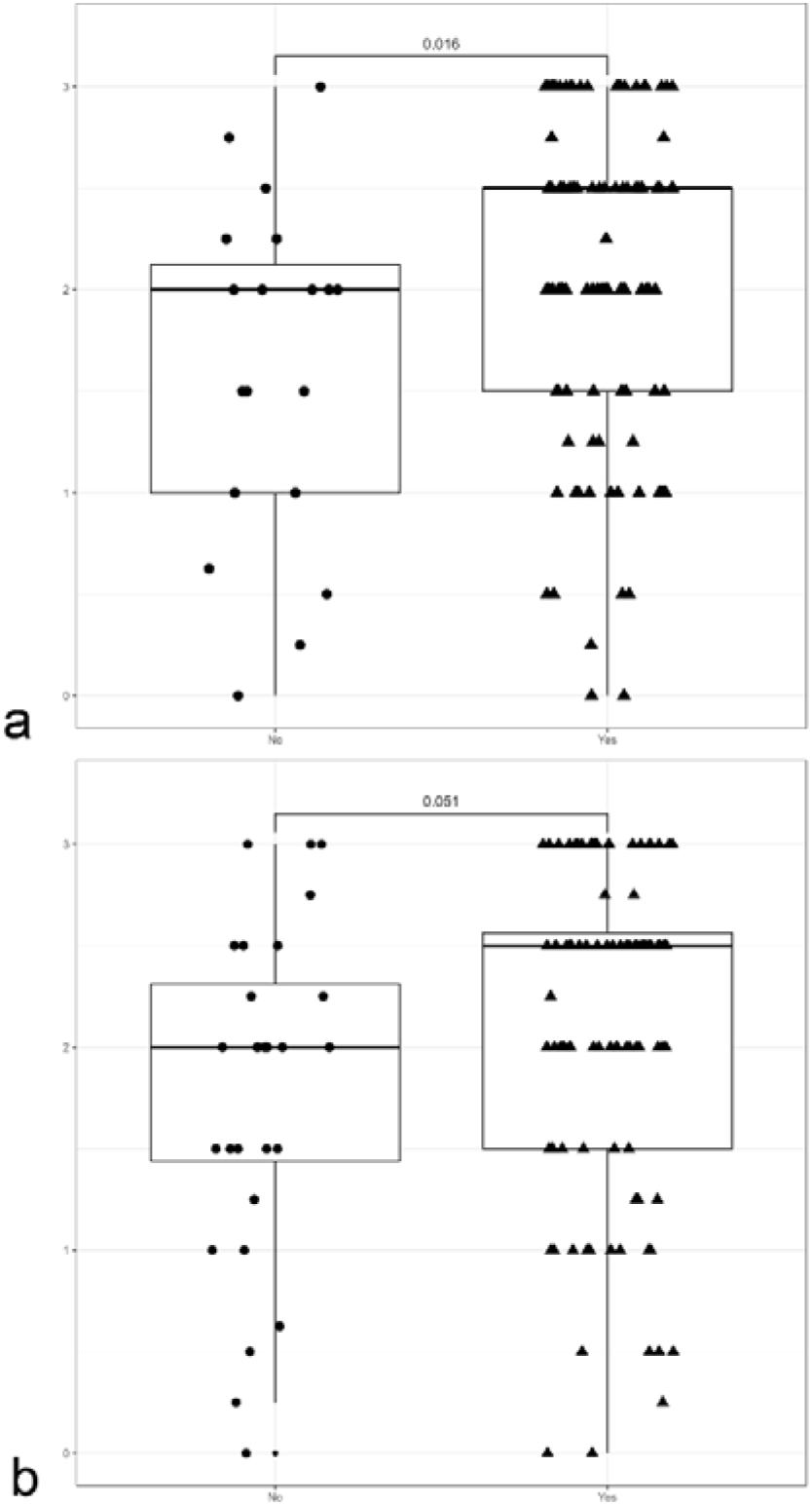
Comparison of mean overall NP expression scores (determined in IHC stained sections) depending on the presence of histopathological changes (**a**) and inflammatory processes (**b**). Only the presence of histopathological changes is associated with significantly higher overall NP scores. No: not present; Yes: present.

We next examined whether inflammatory processes in specific organ systems (enteritis, pneumonia, hepatitis, nephritis) were associated with higher mean overall NP scores. For none of these conditions was there an individual significant association with overall NP scores (Supplemental Figure S5). These findings indicate some association of NP with inflammatory processes in general.

For the spleen we investigated whether the presence of inflammatory processes (splenitis), macrophage rearrangement and granular cell infiltration were associated with overall NP scores and, specifically, with NP scores in the spleen. However, there was no statistically significant association with either (Supplemental Figure S6).

Since several animals exhibited inflammatory processes in more than one organ, we also investigated whether such evidence of systemic inflammatory disease was associated with higher overall NP scores. However, there was no evidence of the latter (Supplemental Figure S7).

### In situ expression of reptarenavirus GP2 and its association with IBs and NP expression

The expression of reptarenavirus NP and the presence of IBs confirms infection in the positive cells, and indicates that replication has occurred or is ongoing. However, efficient completion of virus life cycle, i.e. assembly and egress of virions, requires expression of all viral proteins. We therefore stained consecutive sections for the expression of reptarenavirus GP2, since the glycoproteins are essential for virion formation.^15^ GP2 expression was not consistently detected in the examined tissues (CNS, pancreas, spleen). In fact, we did not detect GP2 in any examined tissues of 12 animals (4 adults, 8 juveniles); however, for most of these, one or two tissues were not available for investigation (Supplemental Table S2). When detected, GP2 expression was often very limited and restricted to rare up to a few individual neurons or ependymal cells in the CNS, individual acini or even individual acinar epithelial cells in the pancreas, and individual leukocytes in the spleen.

Of the 43 brains stained for GP2 expression, 25 (58.14%) harbored at least one positive neuron. In the spinal cord, the percentage was slightly higher (12/18; 66.7%) which was also the case in the pancreas (25/40; 62.5%). The lowest detection rate was found in the spleen (13/38; 34.21%). GP2 expression appeared as a variably intense granular (dot-like) reaction. To specify, in neurons (brain, spinal cord and spinal ganglia), it was represented by a granular reaction that ranged from an individual dot to granular staining of the entire cytoplasm, stretching to or limited to axons, sometimes also as a line of dots along the neuronal cell membrane (Figure 1c). In the pancreas, it was seen in acinar epithelial cells, as a dot like reaction in the cytoplasm or at the luminal surface (Figure 1f). The latter was also seen in ependymal cells in brain and spinal cord (Figure 1c).

While the GP2 expression was too limited or fragmented to be scored, we tried to align it with both the presence of IBs and NP expression and the age of the affected animals.

In most <u>adult animals</u>, GP2 expression was restricted to individual cells in all three tissues, indicating very limited egress of virions (Supplemental Table S2). Nonetheless, GP2 expression was observed in the CNS (12/21 brains), 15/19 pancreata (78.95%), and 8/20 (40%) spleens; for pancreas and spleen the proportion of positive samples was slightly higher than in the entire cohort. However, there were two particularly interesting cases. The first (case 40) was an adult female snake without clinical signs; relevant histological changes were restricted to a mild lymphocyte-dominated gastritis and mixed cellular colitis. The animal exhibited abundant IBs and high NP expression in all examined organs (lung, stomach, intestines, liver, pancreas, spleen, brain, ovary). However, there were also numerous GP2-positive leukocytes, in spleen and the gastric lymphatic tissues; even infiltrating leukocytes in pancreas and kidney were found to express GP2, apparently in the entire cytoplasm (Figure 5b, d, f). These results suggest substantial viral replication in the lymphatic tissues.

**Figure 5.**
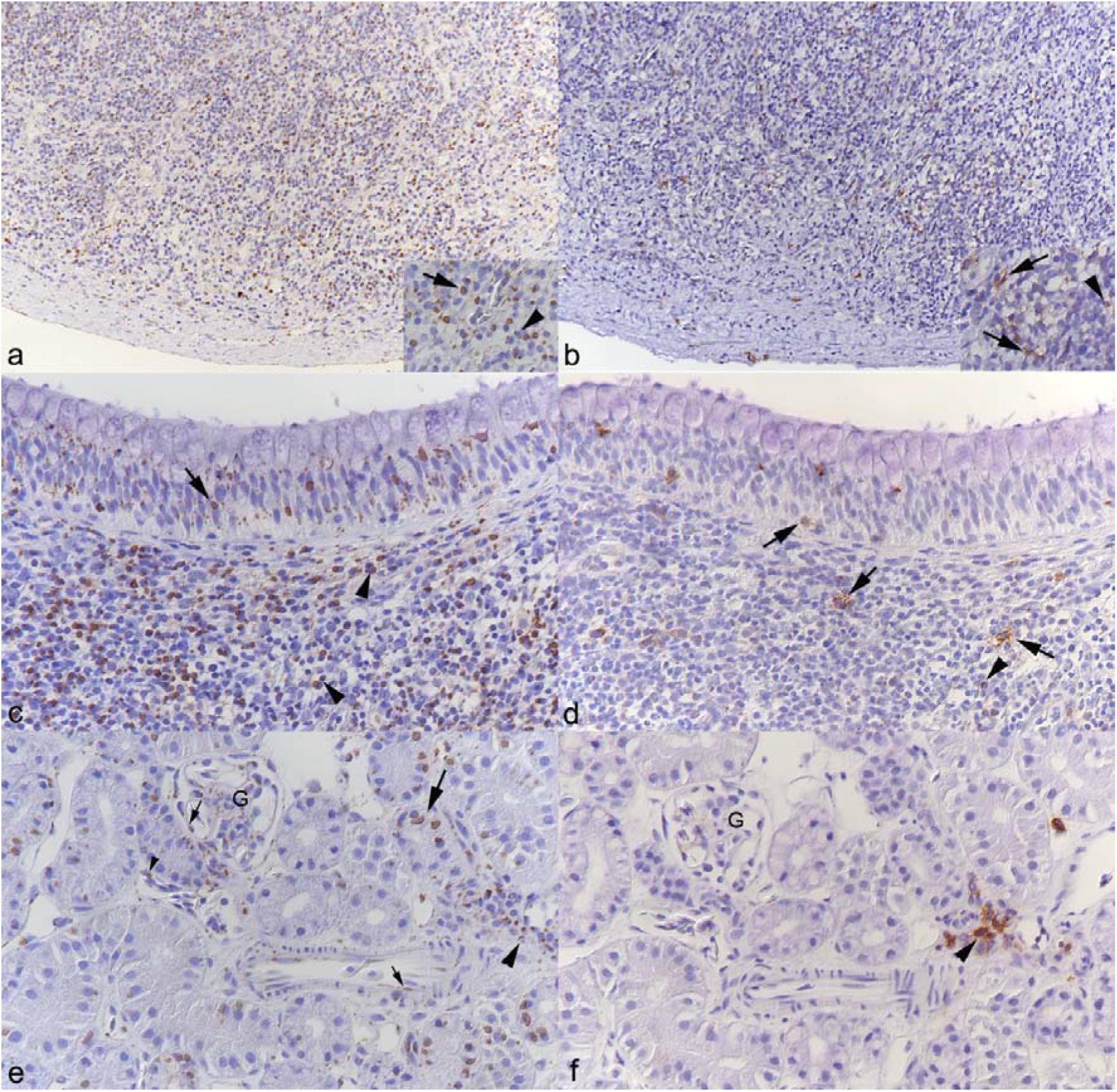
Case 40 (female, 4 years; no clinical signs). Viral protein expression in tissues. Immunohistochemistry (IHC) for viral nucleoprotein (NP; a, c, e) and viral glycoprotein (GP2; b. d, f); hematoxylin counterstain. **a, b)** Spleen. **a)** Most leukocytes exhibit inclusion body (IB) formation with NP expression. Inset: Lymphocytes with small IBs (arrowhead) and large IBs with a weakly positive centre and strongly NP-positive rim (arrow). **b)** Viral GP2 expression is seen in several leukocytes, as granular reaction of the entire cytoplasm (inset: arrows) or punctate cytoplasmic reaction (inset: arrowhead). **c, d)** Stomach. **c)** Viral NP expression is seen in IBs in epithelial cells (arrow) and in leukocytes in the associated lymphatic tissue (arrowheads). **d)** There are scattered GP2-positive leukocytes in the associated lymphatic tissue and between epithelial cells; these show a granular reaction of the entire cytoplasm (arrows) or a punctate cytoplasmic reaction (arrowhead). **e, f)** Kidney. **e)** Viral NP expression is seen in IBs in tubular epithelial cells (large arrow), occasional vascular endothelial cells (small arrows) and intravascular leukocytes (small arrowhead), and in infiltrating leukocytes in the interstitium (large arrowhead). **f)** Infiltrating leukocytes also exhibit strong viral GP2 expression (arrowhead). G: glomerulum.

The second (case 43) was a clinically unremarkable 12-year-old female boa that had been tested positive for BIBD 5 years ago but was kept further by the owner until they could not house it any longer. Euthanasia was decided after unsuccessful rehoming attempts. Histologically, the animal showed mild chronic inflammatory processes in lung, intestine and liver and a focal extensive necrotic hepatitis indicative of bacterial spread from the intestine. The brain exhibited widespread IB formation and NP expression in neurons, and a very mild focal non-suppurative encephalitis, represented by a perivascular infiltrate that contained NP-positive leukocytes (Figure 6a, b). Interestingly, staining for NP also revealed its expression within neurons without overt IB formation (Figure 6c). There were also several strongly GP2-positive neurons and neuronal processes in brain and spinal cord (Figure 6d). These results are consistent with substantial viral replication after or alongside persistence.

**Figure 6.**
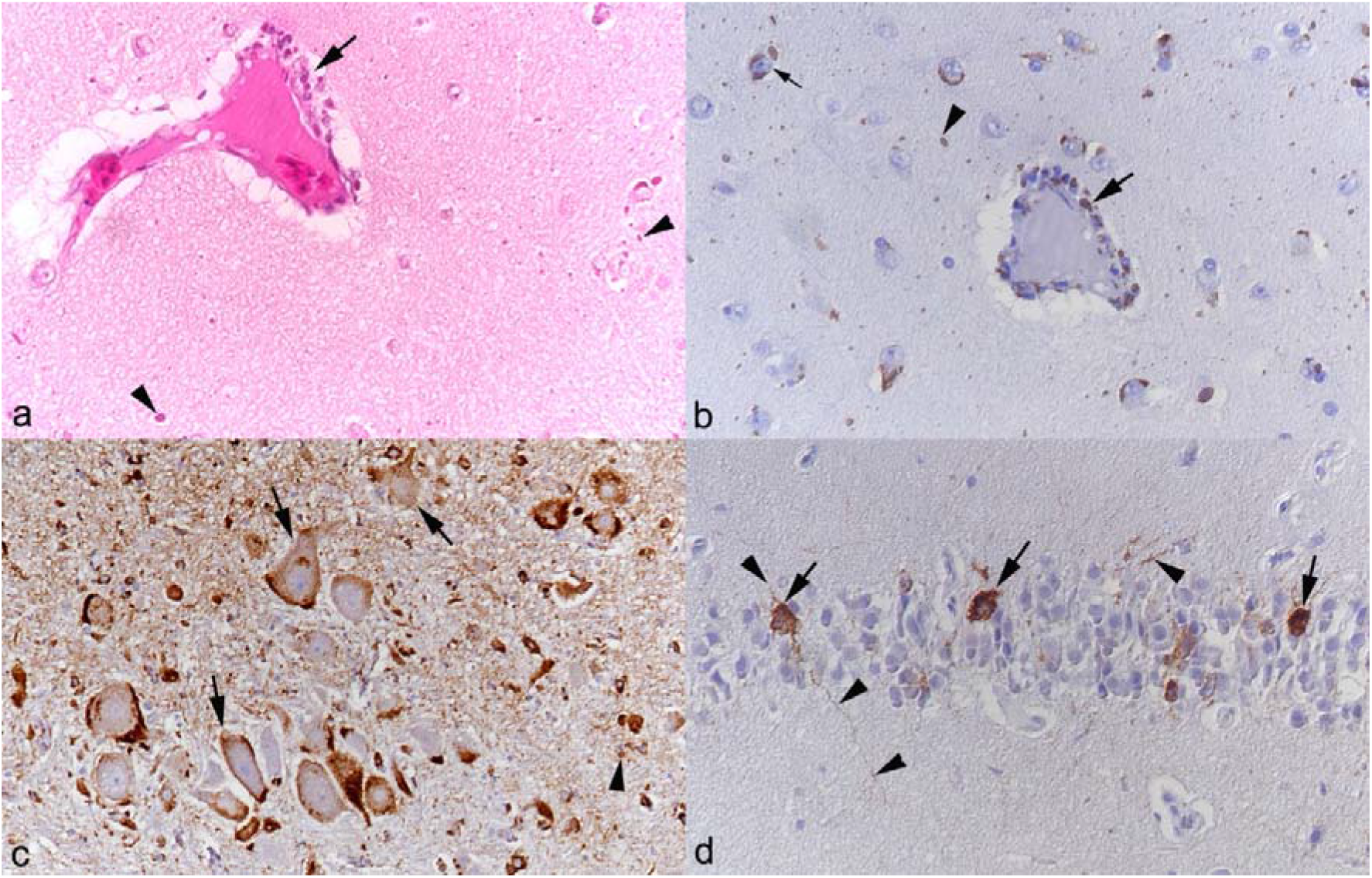
Animal 43 (female, 12 years; no clinical signs). Inclusion body (IB) formation and viral protein expression in the brain. **a, b)** Cortex. **a)** Vessel with mild perivascular leukocyte infiltration (arrow), with IBs in infiltrating leukocytes. HE stain. **b)** Viral nucleoprotein (NP) expression in infiltrating leukocytes (large arrow). There is also evidence of IB formation without clear cellular structures (a, b: arrowheads) and within neurons (b: small arrow). **c)** Olfactory bulb. Numerous large neurons exhibit viral NP expression in the entire cytoplasm, represented by a variably intense granular reaction (arrows). NP is also seen as punctate reactions consistent with IBs (arrowhead). **d)** Medial cortex. Viral glycoprotein (GP2) expression. There is a strong expression in the entire cytoplasm of several neurons (arrows) and expression in neuronal cell processes (arrowheads). b-d) Immunohistochemistry, hematoxylin counterstain.

In most <u>subadult animals</u>, the frequency of GP2 expression in the examined tissues was a bit lower than in the adults (brain and/or spinal cord: 12/23, 52.17%; pancreas: 8/15, 53.3%; spleen: 5/15, 33.3%). However, there were a few animals with more extensive GP2 expression: like their mother (case 40), the 5 siblings (cases 41.1 – 41.5; 4-month-old) exhibited consistent IB and GP2 expression in CNS, pancreas and spleen, some also in bone marrow and/or other lymphatic tissues (cases 41.1, 41.3, 41.4) and each one in stomach and intestine (case 41.1) and liver (case 41.5) which otherwise, like kidney and lung, only harbored IB with NP expression. In a 1-year-old animal (case 26), the brain exhibited very extensive IB formation and NP expression (Figure 7a, d); in some areas, IB appeared to be cell-free; these could represent axons extended by IBs (Figure 7c). Also, occasional neurons showed cytoplasmic vacuolation indicative of degeneration (Figure 7d). These were found to carry IB (and NP; Figure 7e, f) but did not show GP2 expression (Figure 7g). When GP2 expression was observed, it was restricted to individual neurons, and always without IB (Figure 7g, h), presumably representing early stages of infection. Interestingly, we did not detect GP2 expression in the spleen of this animal, suggesting that productive infection does not take place in all organs at the same time.

**Figure 7.**
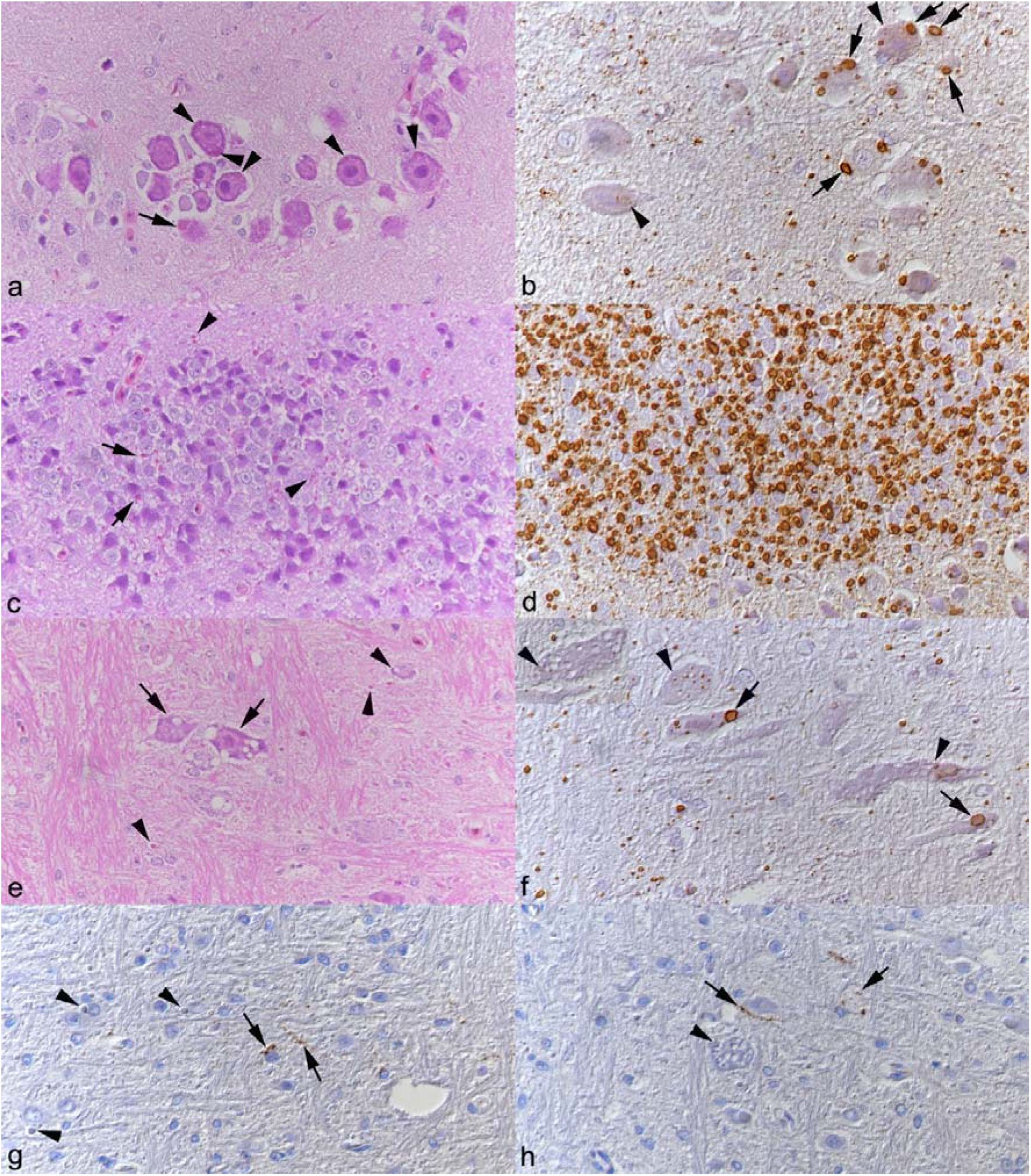
Animal 26 (male, 1 year; no clinical signs). Inclusion body (IB) formation, viral protein expression and histological changes in the brain. **a, b)** Optic tectum (dorsal midbrain). Large neurons carry one to several small IBs (**a**, HE stain). Immunohistochemistry (IHC) for viral nucleoprotein (NP; **b**) highlights these as strongly NP positive (a, b: arrowheads). They are also found to contain large IBs with a weakly NP positive center and a strongly staining rim (a, b: arrows). **c, d)** Cerebellum. Area with very abundant IBs (**c**, HE stain) both in neurons (arrows) and without associated neuronal structures (arrowheads). **d)** IHC for viral NP shows strong expression in IBs. **e, f)** Medulla. **e)** The HE stain highlights a few large neurons with cytoplasmic vacuolation (arrows), indicative of degeneration. The arrowheads point at IBs without neurons, interpreted as IBs inside axons. **f)** Staining for NP by IHC highlights small, strongly NP-positive IBs (arrowheads) also in neurons with cytoplasmic vacuolation (inset). However, there are also neurons that appear morphologically unaltered and contain larger IBs with a weakly stained centre and a rim of pronounced NP expression (arrows). **g, h)** Diencephalon. IHC for viral glycoprotein (GP2). **g)** Several neurons exhibit focal cytopasmic GP2 expression (arrows), whereas others carry distinct IBs (arrowheads) but are GP2 negative. **h)** GP2 expression in the cytoplasm and processes of individual neurons (arrow). Arrowhead: Vacuolated neuron without evidence of GP2 expression. HE stain (a, c, e); IHC; hematoxylin counterstain (b, d, f-h).

In the <u>young animals</u> (neo-/perinatal up to 2 months of age), the brain (and spinal cord) was GP2-positive in 3/4 cases (75%) and the pancreas in 1/4 (25%); the spleen was only available from two and found negative (0/2). However, one neonatal animal (case 14) exhibited very widespread IB formation and viral antigen expression. Specifically, GP2 was expressed in all examined tissues (brain, spinal cord and spinal ganglia, bone marrow and gut associated lymphatic tissue, intestine, pancreas and liver; Figure. 8a-d), in the brain mainly in neurons without IBs, in spinal cord and spinal ganglia also in neurons that carried IBs (which were abundant and often very large; Fig. 8e-g). Interestingly, in another two neonates (cases 15 and 22.2), GP2-positive neurons were more numerous in brain and spinal cord than neurons expressing NP, and when spinal ganglia were available for examination, these also exhibited several GP2-positive neurons. In both animals, bone marrow and thymus were available for examination, and exhibited IBs with NP expression as well as GP2 expression in leukocytes.

**Figure 8.**
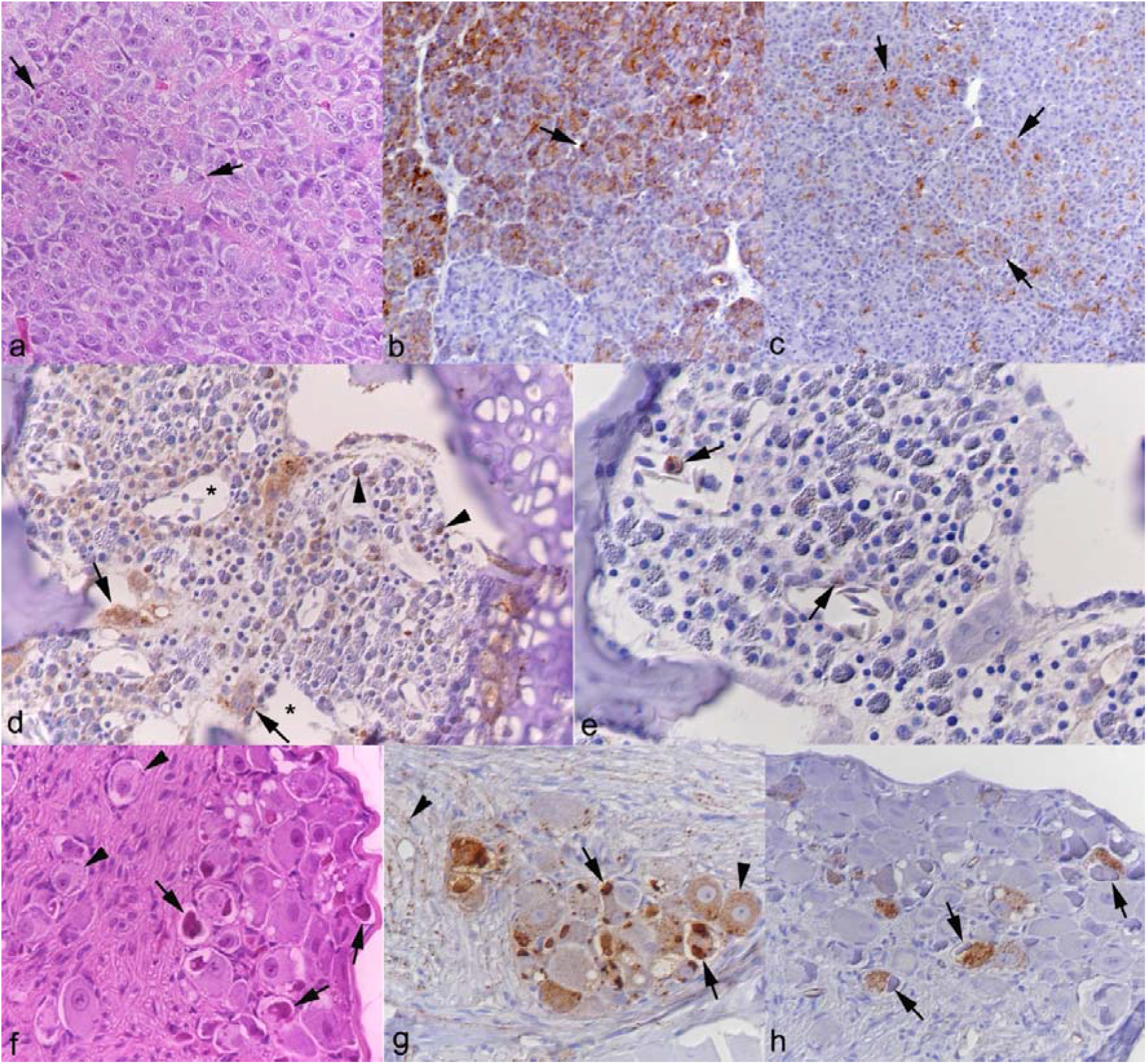
Case 14 (neonate, no clinical signs). Inclusion body (IB) formation and viral protein expression. **a-c)** Pancreas. **a)** The HE stain illustrates the presence of IBs in individual acinar epithelial cells (arrows). **b)** Immunohistology (IHC) for viral NP shows extensive lobular expression of NP in acinar epithelial cells which is occasionally associated with IBs (arrow). **c)** Viral GP2 expression is seen in acinar epithelial cells, mainly in the cytoplasm towards the luminal surface of the cells (arrows). **d, e)** Bone marrow. **d)** Numerous cells exhibit viral NP expression (left). Some cells, close to or around the sinus venosus (asterisks), carry distinct IBs (proerythrocytes,^12^ arrowheads), whereas others show a more finely granular cytoplasmic reaction (arrows). **e)** GP2 expression is seen in individual mononuclear cells with intracytoplasmic granules (progranulocytes;^12^ arows). **f-h)** Spinal ganglion. **f)** The HE stain highlights numerous small round IBs in neurons (arrowheads) as well as several neurons with large, dense, irregularly shaped IBs (arrows). **g)** Staining for viral NP shows its strong expression in the large IBs (arrows) but also in individual small IBs and as a finely granular reaction in the cytoplasm of neurons (arrowheads). **h)** GP2 expression is seen as a granular reaction in neurons; these often carry large IBs (arrows). HE stain (a, f); IHC, hematoxylin counterstain (b-e, g, h).

## Discussion

BIBD is a disease of boid snakes that is caused by reptarenaviruses and morphologically characterized by cytoplasmic inclusion bodies (IB) in most cell types. So far, a concise picture of the clinical and/or pathological features, let alone the pathomechanisms underlying BIBD, was lacking; this was the incentive for the present retrospective study.

We randomly selected a cohort of 50 captive boa constrictors with BIBD, comprising 22 animals from 2000-2003 and 28 from 2013 to 2025. The cohort represented diagnostic case material collected over 26 years in Germany and Switzerland and comprised female and male snakes of different ages (perinatal to 25 years). Animals were systematically assessed for histopathological changes, IB formation and reptarenavirus protein expression.

The earlier cases (n=22; examined between 2000 and 2003) had in the majority shown clinical signs and had been euthanised without or after the diagnosis of BIBD. Reported clinical signs were frequently non-specific (eg. poor body condition, emaciation); when more drastic and specific, these were often related to the CNS, in line with previous reports.^6–10^ Among the 28 cases examined since 2013 were only three that had shown some form of clinical disease, and CNS signs were only reported for a 2-month-old juvenile. This discrepancy could have several reasons; a) increased awareness of snake owners of BIBD; b) *intra vitam* BIBD screening (blood smear based) in collections, prompting euthanasia of affected animals in the attempt to eliminate BIBD from the collection; c) general changes in the disease or the host response over time.

The work-up of the case material provides insight into the pathological changes that occur with BIBD, including those that may predate any clinical disease, since a large proportion of cases were clinically healthy snakes euthanised due to BIBD diagnosis. The vast majority of animals (86%) exhibited pathological changes, albeit often of a low degree. Among these, inflammatory processes were dominating by far. They were detected in 78% (39/50) of the cases and with similar frequency in the old and more recent ones. The alimentary tract was the most commonly inflammation affected organ system, followed by lungs, liver, spleen and kidneys.

The cellular composition varied; however, most represented mixed infiltrates, with macrophage and/or heterophil dominance. In reptiles, both a heterophil-dominated and granulomatous inflammation are generally considered as suggestive of bacterial or fungal infections.^24^ Heterophils and monocyte-derived macrophages, two key cellular components of reptilian innate immunity, can be rapidly recruited to sites of inflammation, with infiltration observed as early as four hours after experimental induction of inflammation in the Brazilian boa (*Boa constrictor constrictor*).^25^ Indeed, in the present study, we detected intralesional bacteria in some cases with granulomatous or fibrinonecrotic inflammation. Notably, animals with enteritis/(entero)colitis frequently showed concurrent inflammatory lesions in additional organs, mainly the liver but also the spleen (sometimes in addition to the liver). This pattern may indicate a compromised intestinal barrier with subsequent bacterial dissemination to extraintestinal sites. Bacterial isolates frequently associated with septicemia and systemic infections in snakes include *Aeromonas* spp., *Pseudomonas* spp., *Salmonella* spp., *Citrobacter* spp., all of which can be part of the normal microbiota and therefore be considered as opportunistic pathogens.^26–28^ The current study did not include bacteriological investigations which is a limitation; therefore, the suspected etiology/pathogenesis could not be verified. Similarly, small lymphoid follicles were observed in the gastric mucosa of jst over 15% of the animals. This finding, which would be termed follicular gastritis in mammals, may reflect chronic antigenic stimulation and is particularly noteworthy given that we have so far observed it exclusively in BIBD-positive animals (personal observation). Together with the inflammatory changes observed in the intestines, this finding supports the interpretation that alimentary tract immunity is altered in the course of BIBD.

In some cases, we observed pneumonia alongside or independent of intestinal inflammatory processes. The pneumonia was generally mild and mostly characterized by lymphocytic infiltrates. No causative infectious agent was identifiable histologically, potentially due to the chronic nature of the lesions, and given the low degree of inflammation, no further diagnostic investigations, such as testing for respiratory viruses, were performed. Nevertheless, the presence of pulmonary inflammation in BIBD-positive snakes suggests that the respiratory tract may represent an additional site of infection.

The current findings are consistent with those of our group’s previous study, in which the histological examination revealed that the majority of BIBD-positive snakes exhibited an enterocolitis of variable severity (23/33 animals; 69%), frequently accompanied by inflammatory lesions in additional organ systems. The study also included a BIBD-negative, clinically healthy *B. constrictor* cohort in which only 11.5% (3/26) exhibited mild inflammatory changes, and limited to a single organ.^12^ Together, both studies provide evidence that BIBD is indeed associated with an increased frequency of inflammatory processes. However, as the organ distribution and histological type of inflammation seem to vary considerably, they are either indicative of bacterial spread or non-specific, and therefore unlikely to represent pathological changes directly attributable to the virus infection.

The high frequency of inflammatory lesions with BIBD is of particular interest considering the longstanding speculation that the disease is associated with immunosuppression in affected snakes,^4,5,9^ a concept that has so far remained largely unsubstantiated by experimental evidence. In a first step to address this question, we recently investigated the immune function of BIBD-positive boa constrictors using *in vitro* assays. This study revealed reduced phagocytic activity of peripheral blood monocytes/azurophils in snakes with BIBD, providing first evidence of impaired function of two phagocytic cell types of reptiles.^29,30^ In mammals, impaired phagocytic function is known to compromise the clearance of invading microorganisms, thereby increasing the risk of secondary infections and prolonging inflammatory responses.^31,32^

Among the 11 animals in which we did not detect any inflammatory processes were the 6 youngest in the study (three neonates/perinatal, each one 1, 2 and 4 months of age). It is likely that these were infected *in utero*, a scenario which we have previously discussed to potentially induce immunological tolerance in the infected offspring.^33^ This phenomenon has been studied in mammarenaviruses, and in particular in murine Lymphocytic choriomeningitis virus infection, in which pre- or neonatal infection can induce immunological tolerance and result in lifelong viral persistence.^34^

The diagnostic hallmark of BIBD are the pathognomonic cytoplasmic IBs that affected animals carry in a broad range of cells including blood cells.^1–3^ Their presence was also the essential inclusion criterion for the present study. The 68 kDa nucleoprotein (NP) of reptarenaviruses is the main component of the IBs,^2–4^ and its detection by IHC confirmed the disease and the underlying viral infection in all cases including those examined prior to the identification of reptarenaviruses as the causative agents of BIBD in 2013.^2,3,14^ Using our large case cohort collected over 26 years we scrutinized the IBs from different qualitative and quantitative angles.

Considering the differences in clinical severity and the general variety of the cases we investigated various parameters after we applied a semiquantitative scoring system for IB abundance in HE stained sections and the extent of NP expression in immunostained sections, respectively, to a defined set of tissues (CNS, pancreas, spleen). Morphometric analysis was not pursued because, e.g., the anatomical regions and amount of tissue available varied among animals in this retrospective case series, limiting the comparability of quantitative measurements across cases. There was no evidence of quantitative differences in association with age or sex. However, the overall scores for both parameters were significantly higher in the more recent material (from 2020 to 2025) compared to the older cases (2000 to 2003). Since this result cannot be explained by prolonged storage of paraffin blocks (which could cause loss of antigenicity), it indicates more extensive IB formation/NP expression in the more recent cohort. Considering that most of these snakes were clinically healthy at euthanasia, this could indicate a decline in virulence of reptarenaviruses over time, allowing for greater virus accumulation in tissues, without disturbance of function. Alternatively, continued breeding of clinically inapparent, virus-positive snakes may have resulted in offspring infected with viral variants exhibiting reduced virulence or different combinations of S and L genome segments.The observation that extensive IB formation and NP expression are not necessarily associated with overt clinical disease is consistent with previous *in vitro* observations, demonstrating that cultured cells can tolerate persistent reptarenavirus infection without significant changes in cell growth or any obvious cytopathic effect, even after repeated passaging.^19,35^ Using the mean overall NP scores for all further assessments, we found these to be lower in the group of animals that had died or were euthanized with clinical signs. This could suggest that more efficient infection in terms of virion production may precede changes leading to clinical disease rather than being responsible for the latter. On the other hand, we found higher mean NP scores with histopathological changes. However, as we observed high variation and a similar spread of scores in both groups, future studies with quantitative approaches are required to investigate this further. Also, delving deeper into the inflammatory processes in different organs (gastrointestinal tract, lung, liver, kidney and spleen) we found no evidence that their presence was associated with a difference in mean NP scores. We also found no evidence of an association with macrophage rearrangement or granular cell infiltration in the spleen, despite the fact that both the latter have so far only been reported in snakes with BIBD.^12^ Similarly, evidence of systemic inflammatory disease, reflected by inflammatory processes in more than one organ, was not associated with higher mean NP scores. All this suggests that these leukocyte-mediated processes are not a direct consequence of virus infection of the tissue and further supports the idea that changes in immune function are responsible for the pathological processes in snakes with BIBD.

In the above context, it is of relevance to gain a clear picture of the effect of reptarenavirus infection in the cells. Accordingly, the histological examination aimed to identify any cellular changes associated with the presence of cytoplasmic IBs, as perceived in HE stained tissue sections, and/or viral NP *in situ* immunolabeling. The most striking finding was the variability of IB size and shape. This was particularly obvious in the CNS, where some neurons harbored very large IBs, occasionally exceeding a diameter of 10 µm, and with an odd shape and a centrally weak NP reaction. An earlier experimental study, where boa constrictors were infected intraperitoneally with a liver homogenate from a BIBD positive boa, described the presence of IBs in liver biopsies collected 10, 20 and 32 weeks post challenge, with progressive increase in number and size of IBs (reaching 5-6 µm at final testing).^4^ Neurons are very long-lived, postmitotic cells; they may indeed progressively accumulate viral protein over time (considering the size, this may indeed be years), whereby the weak NP reaction in the centre of large IBs might be due to a postzone effect where excess antigen prevents sufficient antibody binding.^36^ Evidence of cellular changes in the presence of IBs was extremely limited and restricted to cytoplasmic vacuolation in large neurons in one young snake that had not shown any clinical signs. Indeed, there was no evidence of more (or larger) IBs in the brains of animals with clinical CNS signs; accordingly, virus infection of cells, and the accumulation of NP in the cytoplasmic IB, appears not to be directly cytopathic.^3,19,35^ However, the accumulation of reptarenavirus NP may not be biologically inert.^2–4^ Indeed, taking an *in vitro* approach, our group has previously demonstrated that reptarenavirus NP accumulates in mitochondria.^37^ Although we observed ultrastructural alterations of NP-laden mitochondria in reptarenavirus-infected cells,^37^ their functional impact remains unclear. Whether the observed cytoplasmic vacuolation in some infected large neurons is a consequence of this process, could not be determined. Nonetheless, considering the occurrence of clinical CNS signs without a morphological equivalent apart from the presence of IBs/NP in neurons in several cases, it appears likely that infection can indeed have an impact on cell function. More specifically, the detection of IB in structures consistent with axons raises the question of whether NP accumulation may interfere with neuronal function, particularly neurotransmission. Also, when we investigated the NP expression pattern in cells in more detail, we occasionally detected neurons with a diffuse granular cytoplasmic NP reaction not associated with IBs; these differences in NP organization could indicate recent infection or reactivation of productive infection or virion formation in longer-term infected cells. Interestingly, when we specifically compared the IB abundance and NP expression scores in the tissues in the older and more recent material, we found the mean NP scores in the CNS to be significantly higher than the HE stain based IB scores in the more recent material. Overall, this indicates not only more extensive IB formation but also viral antigen expression and infection of neurons without IB formation in the more recent case material, despite the absence of clinical CNS signs; all this could relate to different S segments or combinations with L segment that differ in the capacity to package or exit the cell efficiently.

In the above context, it is of relevance to also gain more information on viral replication in the tissues. Accordingly, we stained tissue sections consecutive to those stained by HE and for NP for viral glycoprotein (GP2). Notably, GP2 is a structural component of the virion envelope and its marked expression is compatible with ongoing virion assembly and egress, although immunohistology alone cannot prove this.^15,19^ Surprisingly, this expression was far less consistent than the NP expression. Also, while, as expected, it was in principle seen in the same cell types as NP, the expression pattern within cells was different and generally represented by a finely granular cytoplasmic reaction. In leukocytes, this was punctate or throughout the entire cytoplasm, indicating NP accumulation in endoplasmatic reticulum or Golgi apparatus. In contrast, in polarized cells like pancreatic acinar cells and ependymal cells, it was most pronounced at the luminal surface of the cells, possibly suggesting viral release. Neurons showed yet another expression pattern, ranging from a punctate to a diffuse granular reaction in neuronal body and/or axon to punctate reactions along the cell membrane, in neurons that were either free of IBs or contained occasionally even large IBs. The membrane reaction may again represent viral particles ready to be released.^15^ This result indicates that GP2 expression and its intracellular distribution are not necessarily linked to NP formation and accumulation, consistent with a previous observation.^19^ All animals in the current studies were viremic, as they exhibited IBs in blood cells. However, the GP2 staining results suggest that this may not necessarily be associated with permanent substantial virus secretion from leukocytes, since we only detected GP2 expression in leukocytes including precursors in hemolymphatic tissues in a few animals; whether effective systemic spread requires the latter is currently not known. GP2 expression and hence productive infection i.e. virion formation in epithelial cells such as the pancreatic acinar cells suggests virus release and would be consistent with spread with the feces. This was also not consistently observed. Together, these observations point towards infection, i.e. disease, stages or phases, with virus persistence and/or latency, and intermittent recurrent cycles of productive infection. On the other hand, the examination of very young snakes (neonatal to 2 months) revealed substantial GP2 expression in most cells, including a higher proportion of neurons than those expressing NP, indicating an early phase of infection, likely after vertical transmission. This interpretation is consistent with previous *in vitro* observations showing that reptarenavirus GPC and GP2 expression are initially high but decline dramatically during persistent infection, whereas NP expression remains comparatively stable.^19^ Interestingly, although both proteins are encoded by the S segment, GP2 and NP levels appear not to correlate, potentially being regulated differentially. The relatively higher GP2 than NP expression in neurons of very young snakes could also indicate that active viral replication and particle formation during the early stages of infection may favor the incorporation of NP into newly formed virions rather than its intracellular accumulation. Conversely, as hypothesized by Lintala and co-workers, reduced GP expression might impair viral particle formation, leading to NP accumulation and subsequent IB formation.^19^ It is therefore possible that IBs accumulate over time, while viral replication and GP2 expression continue at variable levels.

One animal in the current cohort exhibited a very mild focal nonsuppurative encephalitis with widespread infection and virus replication in neurons, as confirmed by GP2 immunolabeling. The snake had been diagnosed with BIBD 5 years earlier and never shown any clinical signs. This hints at the option of prolonged virus persistence and indicates that extensive viral replication in the brain may induce an inflammatory response, albeit very limited and transient, possibly restricted to phases of particularly high viral load, i.e. replication. It would be tempting to assume that this, together with the inflammatory response, might initiate functional disturbance and, ultimately, clinical signs whereby the latter only become obvious long after any morphological evidence of a tissue response has been resolved. The lack of the latter in the cases with CNS signs in the present study support such an interpretation.

When available from the diagnostic postmortem examination, spinal cord and spinal ganglia were included into the investigation. The latter yielded unexpected results in so far as the largest IBs were found in these neurons. They were also infected in the very young animals with evidence of strong systemic virus replication, and in the neonatal animal where GP2 expression was more intense than NP expression in the brain. This raises the question whether reptarenavirus infection starts from the gastrointestinal tract, with retrograde transneuronal spread from the gastroenteric nerves to the spinal cord and brain, similar to, e.g., pseudorabies virus.^38^ In one adult animal, both NP and GP2 expression was detected in numerous leukocytes in the spleen and gastric lymphatic tissue, as well as in infiltrating leukocytes in pancreas and kidney. This indicates that lymphatic or hematopoetic cells may participate in systemic distribution of the virus and is in line with recent findings in boas with BIBD, where lymphocytes and macrophages in spleen and gut-associated lymphatic tissue, and hematopoietic cells in the bone marrow contained IBs and expressed viral NP.^12^ However, whether such cells actively disseminate virus or only reflect systemic involvement cannot be determined from the present material.

Taken together, the results raise questions regarding the sequence of events of IB formation and productive virus infection which cannot be addressed in naturally infected animals; even experimental infection appears not to be helpful, as so far, there has not been any approach mimicking a potential natural route of infection that has consistently lead to disease in boas.^10,39^ *In vitro* models that mimic the complexity of mixed reptarenavirus infections observed in naturally infected snakes are therefore needed to better understand the dynamics of viral replication, viral protein expression, and IB formation over time.

Some limitations of the study should be considered when interpreting the findings. The retrospective design resulted in some variability in the clinical documentation and tissue collection as part of the diagnostic work-up. However, the number of available samples still allowed meaningful statistical evaluation. A further limitation concerns the semiquantitative nature of IB and NP expression quantification. However, as all samples were examined in direct comparison with each other, their relative correctness can be assumed.

Finally, the time point of infection is not known in our naturally infected case material, and reptarenavirus protein expression and IB formation were assessed histologically and immunohistochemically, without systematic linking to viral RNA levels or viral segment composition in individual animals or tissues. As snakes with BIBD frequently harbor multiple reptarenavirus S and L segments, the lack of systematic segment characterization is particularly relevant because segment composition may influence viral RNA levels, persistence, tissue tropism and host _responses._6,33,40–44

In conclusion, this study shows that BIBD in boa constrictor is associated with frequent inflammatory lesions and widespread reptarenavirus NP expression, but that clinical disease and histological damage cannot be explained simply by the amount of IB or local NP expression. The association between higher overall NP scores and inflammatory lesions supports a link between reptarenavirus protein accumulation and tissue inflammation, while the lack of a consistent local relationship suggests that additional systemic or secondary processes may be involved over time. The morphology and distribution of IBs, together with diffuse NP expression and GP2 expression in selected cells, suggest that different tissues and cells may reflect different phases of reptarenavirus infection. GP2 expression provides a histological indication of ongoing virion assembly or release, but would require confirmation by complementary virological methods, such as quantitative RT-PCR, *in situ* hybridization for viral RNA, detection of replicative RNA intermediates, or virus isolation. Future studies combining standardized tissue sampling, quantitative image analysis and NGS-based segment characterization will be required to further clarify the relationship between viral protein expression, host response and disease progression.

## Acknowledgements

The authors are grateful to the technician teams in the histology laboratories at the Institutes of Veterinary Pathology, University of Giessen, Germany and the Institute of Veterinary Pathology, University of Zurich, for excellent technical support. They also wish to thank the breeders, private owners and reptile shelter for providing animals for this study. We would also like to acknowledge Charlotte Trachsel’s input towards selection, retrieval and initial evaluation of the case material.

