## Supplemental Figures for "Histopathological and virus-associated features of Boid Inclusion Body Disease in *Boa constrictor*"

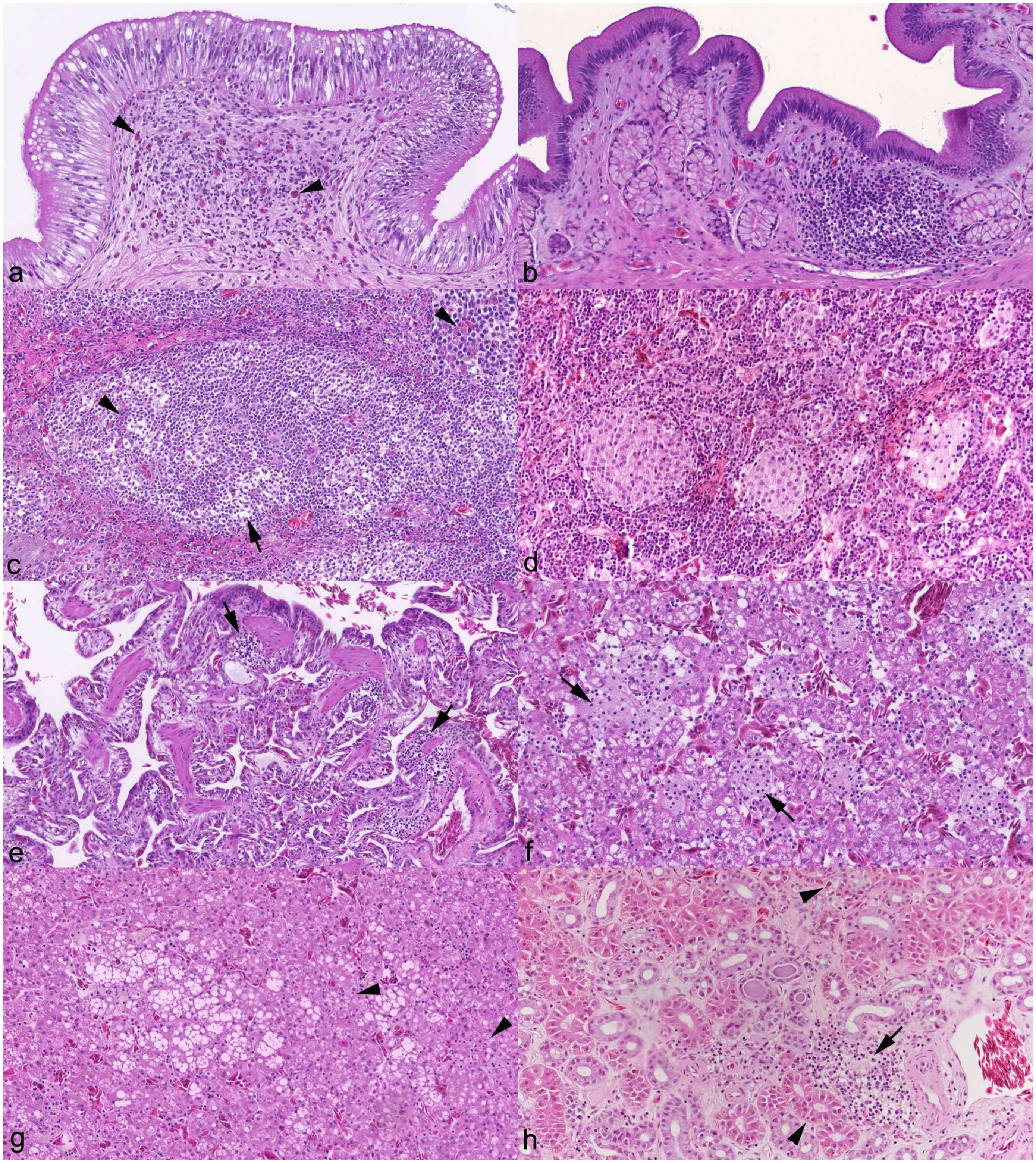


**Supplemental Figure S1**. Representative histopathological changes. **a**. Colon, case 10. Moderate heterophil (arrowheads) dominated colitis. **b**. Stomach, case 40. Mild lymphocyte dominated gastritis. **c.** Spleen, case 23. Focal macrophage rearrangement with presence of granular cells (arrowhead). Arrow: macrophage aggregate. **d**. Spleen, case 10. Multiple granulomas. **e**. Lung, case 17. Multifocal interstitial lymphocyte infiltrates (arrows). **f**. Liver, case 13. Multifocal granulomatous hepatitis. **g**. Liver, case 10. Lipogranulomas. Arrowheads: inclusion bodies within hepatocytes. **h**. Kidney, case 30. Mild focal interstitial lymphocyte infiltration (arrow) and tubular protein casts. Arrowheads: inclusion bodies within tubular epithelial cells.

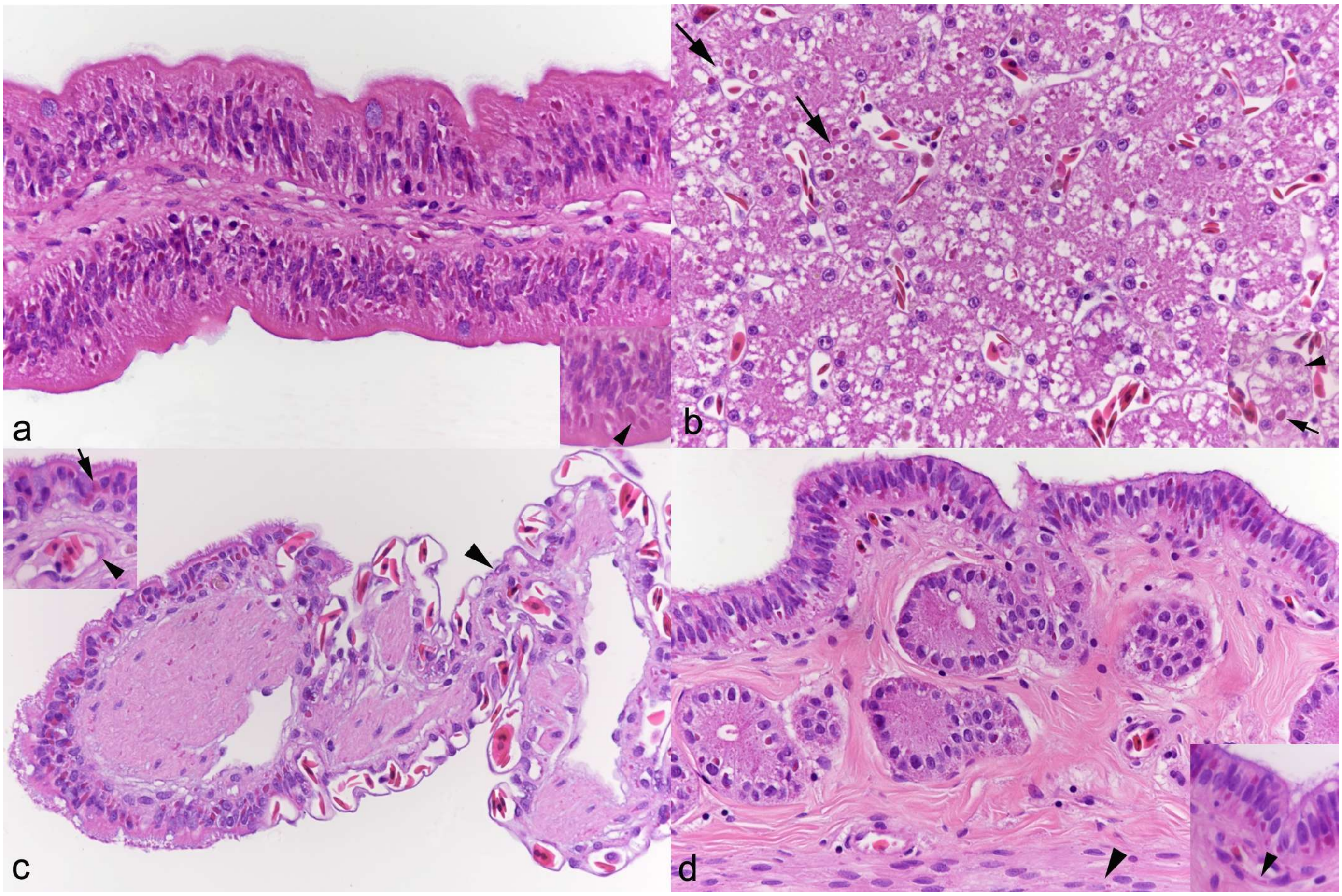


**Supplemental Figure S2**. Illustration of inclusion bodies (IB), based on HE stained sections. Case 40. **a**. Small intestine. Villus. Most enterocytes carry a cytoplasmic IB. Inset: Higher magnification, highlighting enterocytes with one to several (arrowhead) IB. **b**. Liver. Numerous hepatocytes carry IB (arrows). Inset: Higher magnification, highlighting hepatocytes with one large IB (arrow) or two small IB (arrowhead). **c**. Lung. Most ciliated epithelial cells carry IB; rarely, an IB is also apparent in a type I pneumocyte (arrowhead). Inset: Higher magnification, highlighting IB in ciliated epithelial cells (arrow) and in vascular endothelial cells (arrowhead). **d**. Uterus. IB are abundant in endometrial epithelium and glands; small IB are also seen in stromal cells (arrowhead). Inset: Higher magnification, illustrating that IB have a variable position in epithelial cells, at the base and towards the luminal surface. Arrowhead: IB in vascular endothelial cell.

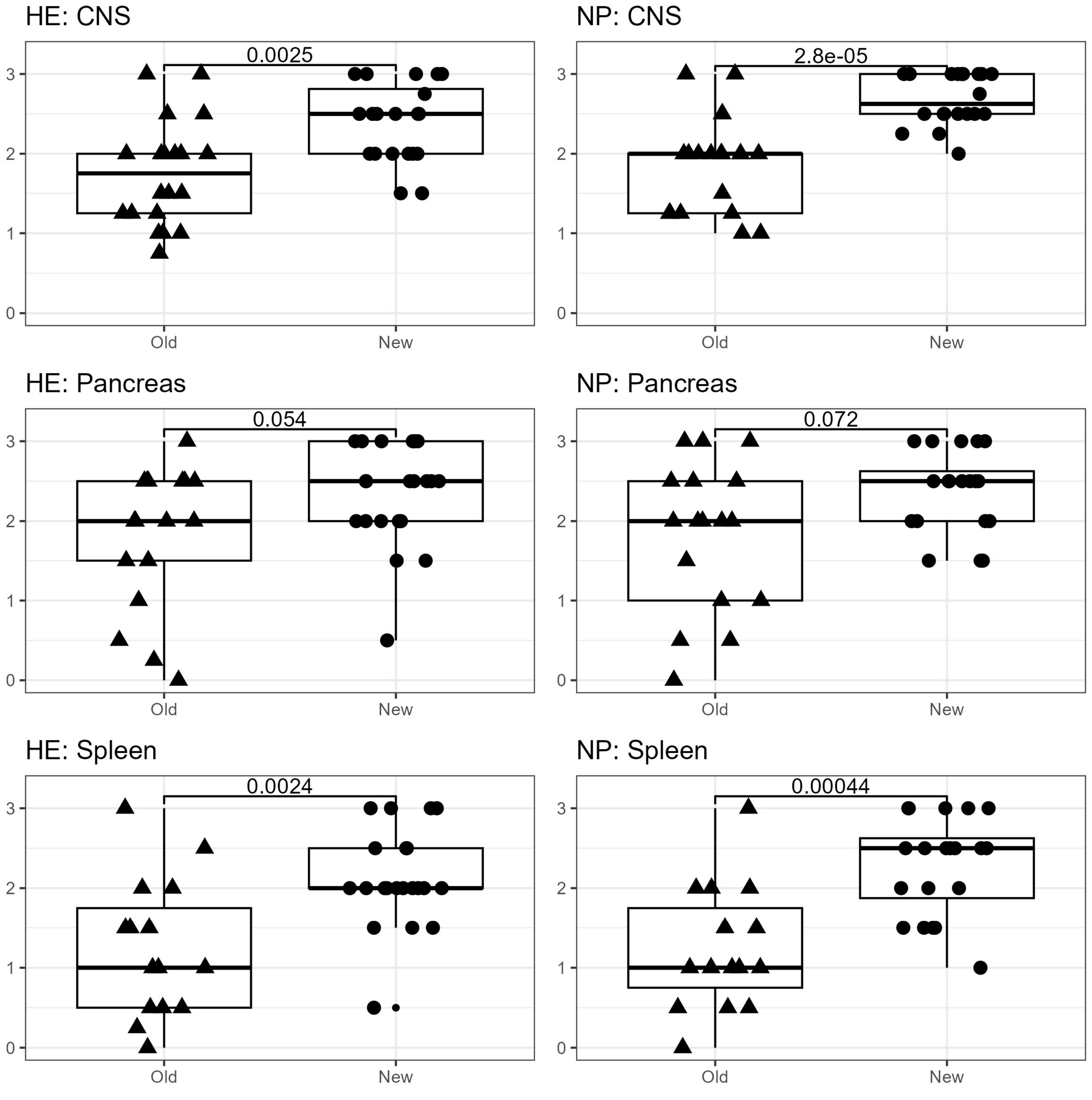


**Supplemental Figure S3**. Comparison of scores in the older archived cases (Old; 2000-2003; n=21) and the more recent cases (New; 2020-2025; n=20) in central nervous system (CNS), pancreas and spleen for inclusion bodies, based on HE stained sections (HE) and based on sections stained for viral nucleoprotein (NP) by immunohistochemistry. In the recent case material, both scores are significantly higher in CNS and spleen.

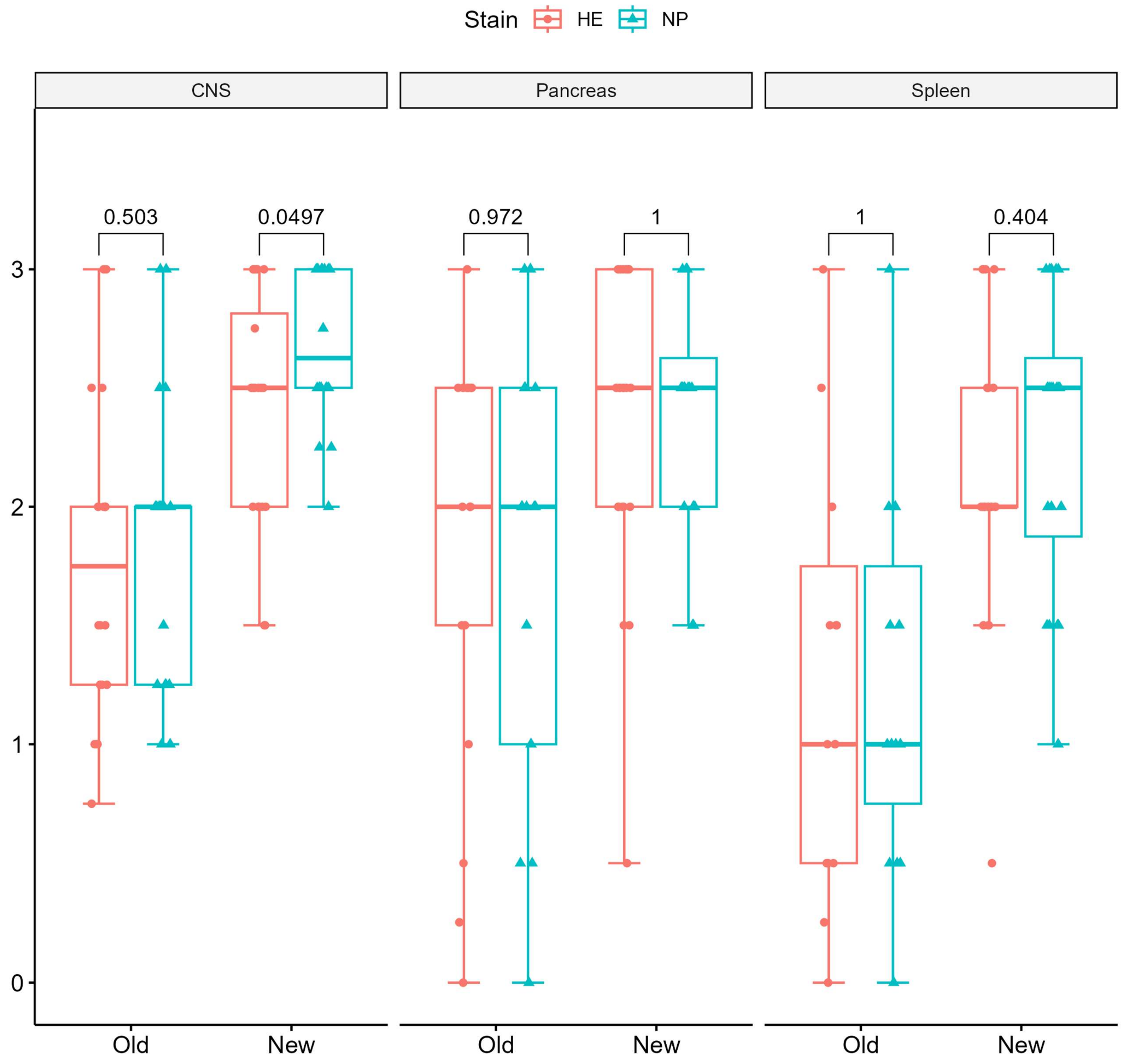


**Supplemental Fig S4**. Comparison of scores for inclusion bodies, based on HE stained sections (HE) and based on sections stained for viral nucleoprotein (NP) by immunohistochemistry, in the older archived cases (Old; 2000-2003; n=21) and the more recent cases (New; 2020-2025; n=20) in central nervous system (CNS), pancreas and spleen. In the recent case material, the NP scores are significantly higher than the HE scores in the CNS but not in pancreas and spleen.

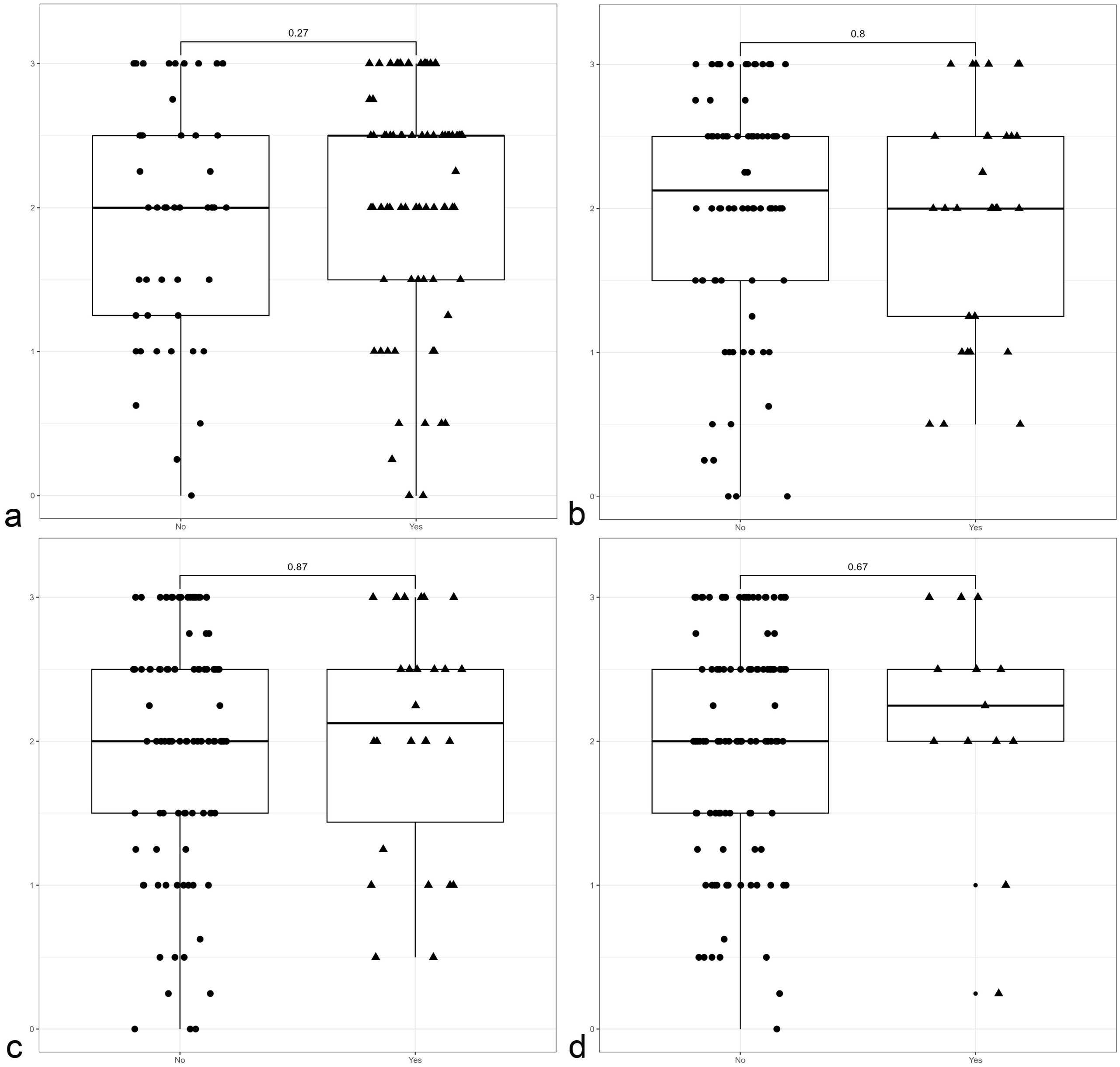


**Supplemental Figure S5**. The extent of reptarenavirus nucleoprotein (NP) expression, based on the scores determined in sections stained for NP by immunohistochemistry, is not associated with the presence of enteritis (a), pneumonia (b), hepatitis (c) or nephritis (d). No: not observed; Yes: observed.

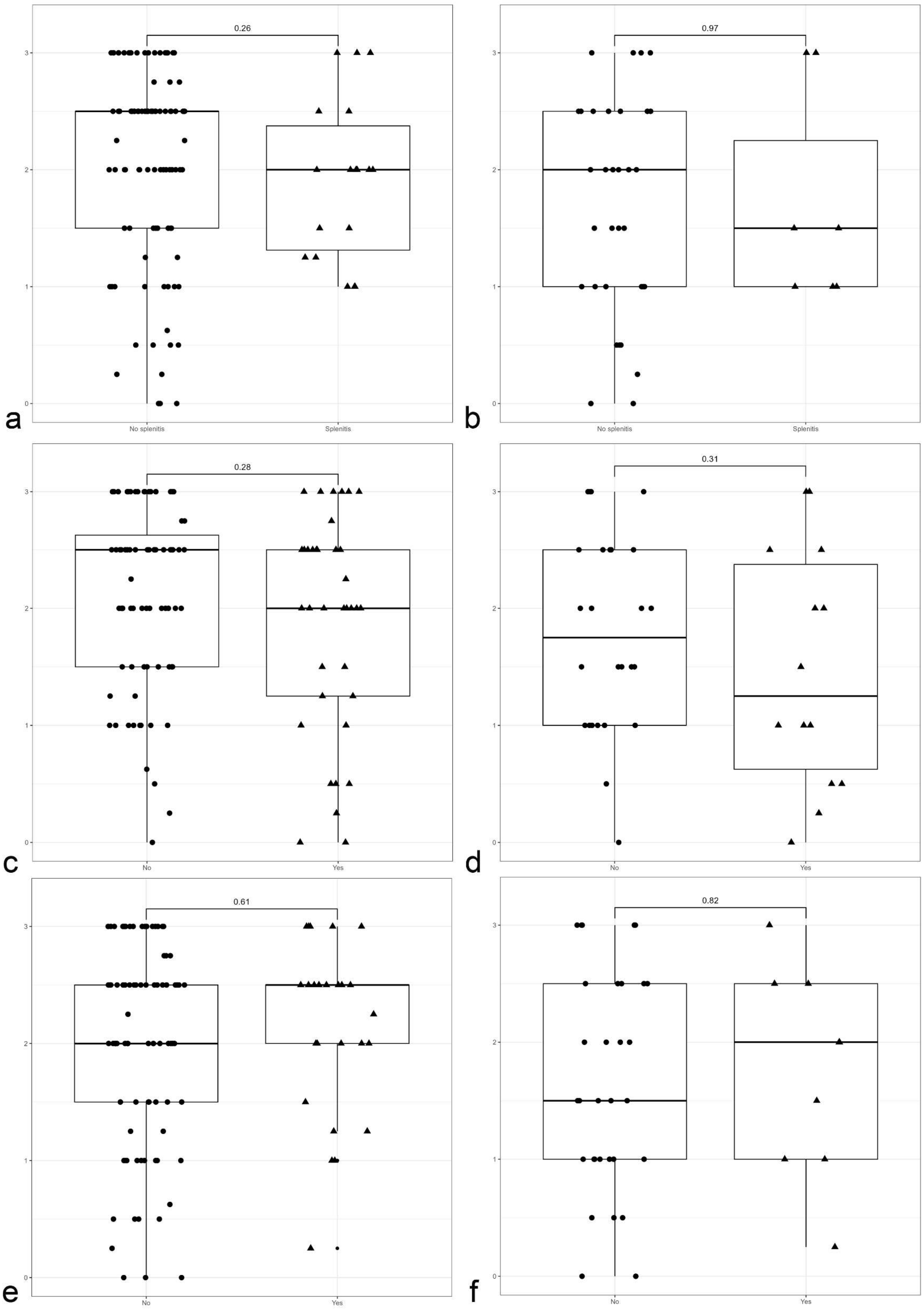


**Supplemental Figure S6**. The extent of reptarenavirus nucleoprotein (NP) expression, based on the scores determined in sections stained for NP by immunohistochemistry, is not associated with the presence of splenitis (a), neither is the NP score in the spleen associated with splenitis (b). The same is true for macrophage rearrangement (c, d) and granular cell infiltration (e, f) in the spleen.

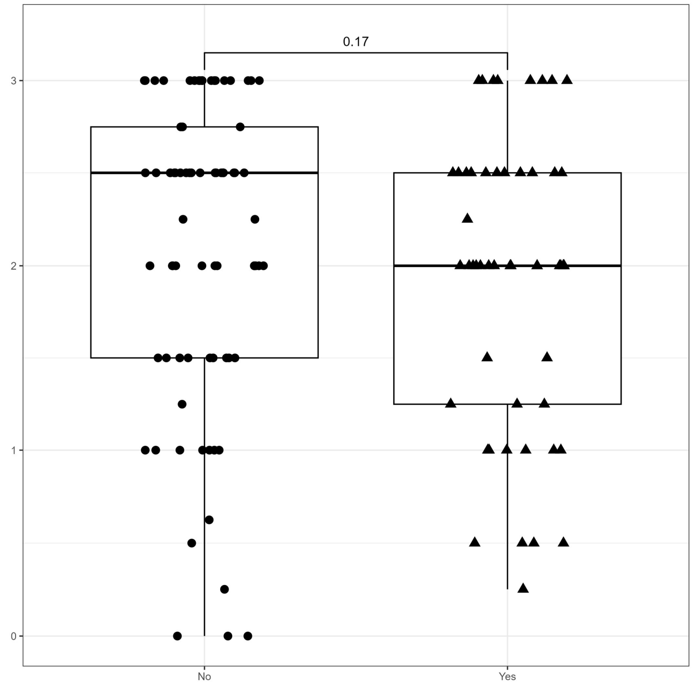


**Supplemental Figure S7**. The extent of reptarenavirus nucleoprotein (NP) expression, based on the scores determined in sections stained for NP by immunohistochemistry, is not associated with the presence of inflammatory processes in more than one organ.
