## Supplemental Table S1 for "Histopathological and virus-associated features of Boid Inclusion Body Disease in *Boa constrictor*"

**Supplemental Table S1**. Animals with confirmed BIBD included into the study, their origin and signalement, with clinical signs and postmortem diagnoses established based on a full diagnostic post mortem examination.

| Animal No | Relationship | Origin[1] | Year[2] | Age[3] | Sex | Size/ Weight | Clinical signs; mode of death | Post mortem diagnoses (organs not available for histological examination) |
|---|---|---|---|---|---|---|---|---|
| 1.1 | Siblings | BF | 2000 | Subadult | NR | 55 g | Poor BC, weakness; euthanasia | Mild diffuse LC-dom enteritis; mild pyogranulomatous hepatitis |
| 1.2 | | | 2000 | Subadult | NR | 65 g | Poor BC, weakness; euthanasia | Mild diffuse heterophil-dom colitis (b) |
| 1.3 | | | 2000 | Subadult | NR | 45 g | Poor BC, weakness, CNS signs; euthanasia | Severe diffuse heterophil- and LC-dom colitis; mild heterophil-dom hepatitis (a, b, e, g) |
| 2 | | BF | 2000 | Subadult | NR | NR | Poor BC, regurgitation, emesis; euthanasia | Mild LC-dom gastritis; mod hepatic lipidosis (a, b, g) |
| 3 | | BF | 2000 | Subadult | NR | NR | Found dead | Mod diffuse heterophil-dom colitis; mild macrophage rearrangement in spleen[4] (h) |
| 4 | | PO | 2000 | Subadult | NR | NR | Poor BC, apathy, CNS signs (headshaking), diarrhea; euthanasia (BIBD)[5] | Severe nematode infection of lung with mild LC-dom interstitial pneumonia; mild LC-dom gastritis; focal ulcerative enteritis; severe granulomatous hepatitis; mod macrophage rearrangement in spleen (b) |
| 5 | | PO | 2001 | Adult | F | 210 cm/ 3.7 kg | Poor BC, anorexia, CNS signs, anisocoria, fibrinonecrotic stomatitis; euthanasia (BIBD) | Severe granulomatous pneumonia with intralesional bacteria; severe granulomatous hepatitis (b) |
| 6 | | PO | 2001 | Subadult | F | 88 g | Poor BC, rhinitis, subcutaneous edema; euthanasia | NHA (a, b, e) |
| 7 | | PO | 2001 | Subadult | NR | NR | Suppurative myositis and osteomyelitis with abscess formation; coprostasis; euthanasia | Suppurative myositis and osteomyelitis with abscess formation; mod diffuse fibrinonecrotic colitis (b, e, g) |
| 8 | | PO | 2001 | Subadult | NR | 80 cm/ 215 g | Poor BC; euthanasia (BIBD) | Mild LC-dom gastritis; mild diffuse LC-dom colitis; mod macrophage rearrangement in spleen (b) |
| 9 | | PO | 2001 | Adult | NR | 135 cm/ 1.34 kg | Poor BC; euthanasia (BIBD) | Mod granular cell infiltration[6] and mod macrophage rearrangement in spleen (b) |
| 10 | | PO | 2001 | 6 y | M | 220 cm/ 4.9 kg | Poor BC, dehydration, CNS signs, respiratory signs, focal fibrinonecrotic stomatitis; euthanasia (BIBD) | Mild multifocal granulomatous enteritis; severe diffuse heterophil- and LC-dom enterocolitis; a few lipogranulomas in liver; mod diffuse interstitial fibrosis in kidneys; mild granulomatous splenitis (b, d, e) |

| | | | | | | | | |
|---|---|---|---|---|---|---|---|---|
| 11 | | PO | 2001 | Subadult | M | 106 cm/ 580 g | Severe emaciation; euthanasia (BIBD) | Multiple liver abscesses with intralesional bacteria (b) |
| 12 | | PO | 2001 | Adult | F | 189 cm/ 5.5 kg | Euthanasia (BIBD) | Mild LC-dom gastritis; severe multifocal pyogranulomatous colitis with intralesional bacteria; mild heterophil-dom hepatitis; severe granulomatous salpingitis with intralesional bacteria (a, b, g) |
| 13 | | BF | 2002 | 4 y | F | 147 cm/ 714 g | Abscess Jacobson's (vomeronasal) organ, myositis and osteomyelitis; euthanasia (BIBD) | Pyogranulomatous myositis and osteomyelitis; mod diffuse heterophil-dom colitis; mod granulomatous hepatitis; mild granular cell infiltration in spleen; severe granulomatous splenitis (c, d, e) |
| 14 | | BF | 2003 | Neonate | NR | NR | Euthanasia (BIBD) | NHA (a, b) |
| 15 | | BF | 2003 | Neonate | NR | NR | Coelomic hernia; euthanasia | NHA (a) |
| 16 | | PO | 2003 | Subadult | NR | NR | Euthanasia (no further information available) | Mild multifocal LC-dom interstitial pneumonia; mild granular cell infiltration and mod macrophage rearrangement in spleen; mod granulomatous splenitis; epithelial cysts in thymus (e) |
| 17 | Father of Animal 19 | PO | 2003 | Adult | M | NR | Euthanasia (BIBD) | Mod LC-dom interstitial pneumonia; mod diffuse heterophil-dom colitis (a, e, f, g) |
| 18 | Mother of Animal 19 | PO | 2003 | Adult | F | NR | Euthanasia (BIBD) | Mild chronic heterophil- and LC-dominated interstitial pneumonia; severe granular cell infiltration and macrophage rearrangement in spleen; mild granulomatous splenitis (b, e) |
| 19 | Offspring of Animals 17 and 18 | PO | 2003 | 6 mo | NR | NR | Euthanasia (BIBD) | Mild diffuse heterophil-dom colitis; mod chronic LC-dom interstitial pneumonia; severe macrophage rearrangement in spleen (b, c) |
| 20 | | PO | 2003 | Adult | F | NR | Euthanasia (BIBD) | Mod diffuse heterophil-dom colitis (b, d, e) |
| 21 | | PO | 2013 | 1 mo | NR | NR | Euthanasia (BIBD) | Moderate multifocal hepatic lipidosis and degeneration (a, b) |
| 22.1 | Siblings | BF | 2014 | 2 mo | NR | 55 g | Lack of righting reflex; euthanasia (BIBD) | NHA (b) |
| 22.2 | | | 2014 | Perinatal | NR | NR | Euthanasia (BIBD) | NHA |
| 23 | | BF | 2017 | Adult | F | NR | Euthanasia (BIBD) | Focal cholesterol cleft granuloma in lung; mild diffuse heterophil- and LC-dom colitis; |

| | | | | | | | | |
|---|---|---|---|---|---|---|---|---|
| | | | | | | | | mild LC-dom pyelonephritis; severe macrophage rearrangement and severe granular cell infiltration in spleen (b) |
| 24 | | PO | 2018 | Adult | M | 162 cm/ 960 g | Anorexia, emaciation, rhinitis; death | Severe diffuse fibrinonecrotic enteritis; mild necrotic hepatitis with intralesional bacteria; mild interstitial LC-dom nephritis; mod granulomatous and necrotic splenitis with intralesional bacteria and vascular thrombi (b, e, g) |
| 25 | | BF | 2019 | 7 y | M | NR | Euthanasia (BIBD) | Mild granulomatous gastritis; severe macrophage rearrangement in spleen (b) |
| 26 | | BF | 2020 | 1 y | M | 83 cm/ 350 g | Euthanasia (BIBD) | Mild multifocal heterophil-dom enteritis; mild focal granulomatous hepatitis |
| 27 | | BF | 2020 | 1 y | M | 75 cm/ 300 g | Euthanasia (BIBD) | Mild diffuse heterophil-dom pneumonia; mild diffuse heterophil-dom enteritis |
| 28 | | BF | 2020 | 2 y | F | 120 cm/ 1.62 kg | Euthanasia (BIBD) | Mild diffuse heterophil-dom colitis |
| 29 | | BF | 2020 | 2 y | F | 140 cm/ 2.6 kg | Euthanasia (BIBD) | Mod diffuse heterophil-dom colitis (b, d) |
| 30 | | BF | 2020 | 8 y | M | 90 cm/ 800 g | Emaciation; euthanasia (BIBD) | Mod diffuse heterophil- and LC-dom interstitial pneumonia; mild diffuse heterophil-dom enterocolitis; mild LC-dom interstitial nephritis (b) |
| 31 | | BF | 2020 | 10 y | F | 270 cm/ 10.7 kg | Euthanasia (BIBD) | Severe diffuse heterophil-dom colitis; moderate bile stasis (e) |
| 32 | | BF | 2020 | 11 y | F | 232 cm/ 10.1 kg | Euthanasia (BIBD) | Mild diffuse LC-dom tracheitis, interstitial pneumonia and hepatitis |
| 33 | | BF | 2020 | 11 y | F | 250 cm/ 10.9 kg | Euthanasia (BIBD) | Mild diffuse LC-dom interstitial pneumonia; mild diffuse heterophil-dom colitis; nodular hyperplasia of exocrine pancreas; mod heterophil-dom nephritis with focal dystrophic mineralization |
| 34 | | BF | 2020 | 5 y | M | 185 cm/ 3.3 kg | Euthanasia (BIBD) | Mild heterophil-dom splenitis; mild macrophage rearrangement in spleen (g) |
| 35 | | BF | 2020 | 3 y | M | 213 cm/ 1.7 kg | Euthanasia (BIBD) | Mild diffuse heterophil and LC-dom interstitial pneumonia (b) |
| 36 | | BF | 2020 | Adult | F | 213 cm/ 7.2 kg | Euthanasia (BIBD) | Mod diffuse heterophil-dom colitis; mild heterophil-dom splenitis (d) |

| | | | | | | | | |
|---|---|---|---|---|---|---|---|---|
| 37 | | RC | 2020 | 25 y | F | NR | Euthanasia (BIBD) | Mod diffuse hepatic lipidosis; mild macrophage rearrangement in spleen; epithelial cysts in thymus |
| 38 | | PO | 2022 | 13 y | F | 170 cm/ 5.3 kg | Euthanasia (BIBD) | Mod diffuse heterophil-dom enterocolitis (b, c) |
| 39 | | PO | 2022 | 13 y | M | 140 cm/ 7.6 kg | Euthanasia (BIBD) | NHA (c) |
| 40 | Mother of 41 | RC | 2024 | 4 y | F | 220 cm/ 5.2 kg | Euthanasia after several unsuccessful rehoming attempts | Mild LC-dominated gastritis; mild diffuse heterophil- and LC-dom colitis; mild macrophage rearrangement and mild granular cell infiltration in spleen; epithelial cysts in thymus |
| 41.1 | Siblings, offspring of 40 | RC | 2024 | 4 mo | F | NR | Euthanasia due to BIBD positive mother (animal 40) | Severe diffuse heterophil- and LC-dom and granulomatous colitis (b) |
| 41.2 | | | 2024 | 4 mo | M | NR | | Severe diffuse heterophil- and LC-dom colitis; epithelial cysts in thymus |
| 41.3 | | | 2024 | 4 mo | M | NR | | Mod diffuse heterophil-dom colitis; mild granular cell infiltration and macrophage rearrangement in spleen |
| 41.4 | | | 2024 | 4 mo | F | NR | | NHA (b) |
| 41.5 | | | 2024 | 4 mo | M | NR | | Mild diffuse heterophil-dom colitis (a, b) |
| 42 | | RC | 2024 | Adult | M | NR | Euthanasia (BIBD) | Mild granular cell infiltration in spleen |
| 43 | | PO | 2025 | 12 y | F | NR | Tested positive for BIBD 5 y ago; euthanasia after unsuccessful rehoming attempts | Mild multifocal LC-dom interstitial pneumonia; mild diffuse heterophil-dom colitis; focal extensive necrotic hepatitis; mod LC-dom interstitial nephritis with fibrosis; mild macrophage rearrangement and mild granular cell infiltration in spleen. |

Abbreviations: BC – body condition; dom – dominated; F – female; LC – lymphocyte; M – male; mo – month(s); mod – moderate; NHA – no histological abnormalities; NR – not recorded; y – year(s)

1 Origin: This provides information whether the animals originated from a breeding facility (BF), with knowledge on group size and composition as well as relationship, were privately owned (PO) individually kept animals without information on group size and composition, or were housed in a rescue center (RC), i.e. a shelter, without information on their background.

2 This indicates the year of death, post mortem examination and tissue sampling.

3 Subadult: <3 years of age, i.e. prior to definite sexual maturity and reproduction; adult: ≥3 years of age, i.e. sexually mature, reproducing animals (as described in [22]).

4 Aggregates of macrophages consistent with a granulomatous response (as described in [12]).
5 The wording "euthanasia (BIBD)" implies that the animal was euthanized due to confirmed BIBD.
6 Presumed leukocytes of unknown lineage that increase in number with advancing age (as described in [21]).

Organs not available for histological examination in individual animals:
a. Spleen
b. Thymus
c. CNS (brain and spinal cord)
d. Lung
e. Stomach
f. Liver
g. Pancreas
h. Kidney
