## Supplemental Table S2 for "Histopathological and virus-associated features of Boid Inclusion Body Disease in *Boa constrictor*"

**Supplemental Table S2.** Results of the semiquantitative assessment of the presence of cytoplasmic inclusion bodies determined in HE stained sections (HE), and reptarenavirus nucleoprotein (NP) expression determined in sections stained by IHC, in cells in brain, spinal cord (incl. spinal ganglia), pancreas and spleen of boa constrictors with BIBD (n=50; for details, see Supplemental Table S1). The presence and extent of the viral glycoprotein 2 (GP2) expression, as an indicator of viral replication, was also determined in sections stained by IHC and is commented on specifically for each case where expression was observed.

| Animal No | Brain | | | Spinal cord (+ ggl.) | | | Pancreas | | | Spleen | | | Comment (GP2 expression) |
|---|---|---|---|---|---|---|---|---|---|---|---|---|---|
| | HE | NP | GP2 | HE | NP | GP2 | HE | NP | GP2 | HE | NP | GP2 | |
| 1.1 | NE | NE | NE | 2 | 2.5 | + | 2.5 | 2.5 | - | 0.5 | 0.5 | - | SC: dot-like reaction along cell periphery (neurons incl. axons, ependymal cells) |
| 1.2 | NE | NE | NE | 2 | 2 | (+) | 2 | 2 | - | 1 | 1 | - | SC: very rare ind pos neurons |
| 1.3 | 1 | 1 | - | 0.5 | 1 | - | NE | NE | NE | NE | NE | NE | |
| 2 | 2 | 2 | - | 1 | 1 | - | NE | NE | NE | NE | NE | NE | |
| 3 | 2 | 2 | - | NE | NE | NE | 0 | 0 | - | 0 | 0 | - | |
| 4 | 2.5 | 2.5 | - | 1.5 | 1.5 | - | 2.5 | 2.5 | - | 0.5 | 0.5 | - | |
| 5 | 2 | 2 | - | 0.5 | 0.5 | - | 1.5 | 1 | - | 1.5 | 1 | - | |
| 6 | 2 | 2 | - | NE | NE | NE | 2 | 2 | - | NE | NE | NE | |
| 7 | 2.5 | 2 | - | NE | NE | NE | NE | NE | NE | 2.5 | 2 | - | |
| 8 | 3 | 3 | + | NE | NE | NE | 2.5 | 2.5 | + | 1.5 | 1.5 | - | Brain: dot-like reaction along cell periphery (neurons incl. axons) in ind cells; pancreas: scattered acini with reaction along luminal surface of AEC |
| 9 | 1.5 | 1.5 | (+) | 1 | 1 | (+) | 2.5 | 3 | (+) | 1.5 | 2 | (+) | Brain, SC: dot-like reaction along cell periphery (neurons incl. axons) of very rare ind cells; spleen: very rare ind leukocytes; pancreas: weak reaction in rare ind AEC |
| 10 | 2 | 2 | + | 0.5 | 0.5 | - | 2.5 | 2 | + | 2 | 1.5 | (+) | Brain: random ind neurons with cytoplasmic and/or peripheral reaction; spleen: rare ind leukocytes; pancreas: mod number of random ind AEC |
| 11 | 2.5 | 2.5 | - | NE | NE | NE | 3 | 3 | - | 2 | 2 | - | |
| 12 | 1.5 | 2 | - | NE | NE | NE | NE | NE | NE | NE | NE | NE | |
| 13 | NE | NE | NE | NE | NE | NE | 2.5 | 3 | - | 3 | 3 | - | |

| | | | | | | | | | | | | | |
|---|---|---|---|---|---|---|---|---|---|---|---|---|---|
| 14 | 3 | 3 | + | 3 | 3 | + | 1 | 1 | + | NE | NE | NE | Brain: expression mainly in neurons without IB; SC, SG: expression also in neurons with IB; pancreas: numerous AEC |
| 16 | 2 | 2 | - | NE | NE | NE | 2 | 2 | - | 1 | 1 | - | |
| 15 | 2 | 2 | + | 1 | 2 | + | 0.5 | 0.5 | - | NE | NE | NE | Brain, SC: more numerous pos neurons than NP-pos neurons |
| 17 | 1 | 2 | + | NE | NE | NE | NE | NE | NE | NE | NE | NE | Brain: random ind neurons with cytoplasmic and/or peripheral reaction |
| 18 | 1 | 1 | (1) | 1 | 1.5 | (+) | 2 | 2 | - | 1 | 1 | - | Brain: dot-like reaction along cell periphery (neurons incl. axons) in ind cells; SC: two ependymal cells with expression at luminal surface |
| 19 | NE | NE | NE | NE | NE | NE | 0.25 | 0.5 | + | 0.25 | 0.5 | + | Pancreas: numerous acini with positive AEC; spleen: scattered ind leukocytes |
| 20 | 1 | 1 | + | 1 | 1 | (+) | 1.5 | 1.5 | (+) | 0.5 | 1 | - | Brain: dot-like reaction along cell periphery (neurons incl. axons) in some ind cells; SC: weak reaction in rare ind neuron and ependymal cell; pancreas: scattered ind AEC |
| 21 | 2 | 2.5 | (+) | 1 | 2 | + | 1 | 2 | - | NE | NE | NE | Brain: very rare ind neuron; SC: several ind neurons with dot-like cytoplasmic reactions |
| 22.1 | 2.5 | 3 | - | 2.5 | 2.5 | NE | 1.5 | 2 | - | 0 | 1 | - | |
| 22.2 | 0.25 | 0.25 | + | 1 | 1 | + | 0.25 | 0.25 | - | 0 | 0 | - | Brain: more numerous pos neurons than NP-pos neurons; SC, SG: several pos neurons |
| 23 | 3 | 3 | - | NE | NE | NE | 3 | 2.5 | + | 2 | 0.25 | (+) | Pancreas: random acini with pos AEC; spleen: scattered ind pos leukocytes |
| 24 | 2 | 2 | - | NE | NE | NE | NE | NE | NE | 1.5 | 1 | (+) | Spleen: scattered ind pos leukocytes |
| 25 | 1.75 | 2.75 | + | NE | NE | NE | 2.5 | 2.5 | (+) | 2 | 1 | (+) | Brain: numerous neurons; pancreas: scattered ind AEC; spleen: scattered ind leukocytes |
| 26 | 3 | 3 | + | NE | NE | NE | 2 | 2.5 | (+) | 1.5 | 2 | - | Brain: several neurons and axons, mainly with dot-like reaction along cell periphery; pancreas: rare ind AEC |
| 27 | 2 | 2.5 | (+) | NE | NE | NE | 1.5 | 2 | - | 1.5 | 2.5 | - | Brain: very few neurons and axons with dot-like reaction along cell periphery |
| 28 | 1.5 | 2.5 | (+) | NE | NE | NE | 2 | 2.5 | - | 2 | 3 | - | Brain: one pos neuron |

| | | | | | | | | | | | | | |
|---|---|---|---|---|---|---|---|---|---|---|---|---|---|
| 29 | 2 | 3 | (+) | NE | NE | NE | 2.5 | 2.5 | + | 2.5 | 3 | - | Brain: scattered ind neurons; pancreas: random acini with pos AEC |
| 30 | 3 | 3 | + | NE | NE | NE | 2 | 2 | + | 3 | 2.5 | - | Brain: ind neurons and processes; pancreas: random acini with pos AEC |
| 31 | 2.5 | 3 | - | NE | NE | NE | 2.5 | 2.5 | (+) | 2 | NE | NE | Pancreas: scattered ind AEC |
| 32 | 2.5 | 3 | - | NE | NE | NE | 3 | 3 | + | 2.5 | 2.5 | - | Pancreas: several random acini (AEC) and scattered ind AEC |
| 33 | 3 | 3 | - | NE | NE | NE | 2 | 2 | + | 2.5 | 2.5 | NE | Pancreas: several random pos acini (AEC) |
| 34 | 2 | 2.5 | - | NE | NE | NE | NE | NE | NE | 3 | 3 | - | |
| 35 | 3 | 3 | - | NE | NE | NE | 3 | 3 | (+) | 3 | 3 | - | Pancreas: rare ind AEC |
| 36 | 2 | 2.5 | (+) | NE | NE | NE | 2 | 2 | (+) | 1.5 | 1.5 | - | Brain: one pos neuron; pancreas: scattered ind AEC |
| 37 | 1.5 | 2 | (+) | NE | NE | NE | 1.5 | 1.5 | + | 3 | 3 | - | Brain: rare ind pos neurons; pancreas: random pos acini (AEC) and ind AEC |
| 38 | NE | NE | NE | NE | NE | NE | 3 | 1.5 | (+) | 2 | 1 | - | Pancreas: very rare ind AEC |
| 39 | NE | NE | NE | NE | NE | NE | 2.5 | 1.5 | (+) | 2 | 1.5 | - | Pancreas: scattered ind AEC |
| 40 | 2.5 | 2.5 | + | NE | NE | NE | 2.5 | 2.5 | + | 2 | 2.5 | + | Brain: numerous ind neurons and axons; pancreas: a few random acini with pos AEC; spleen: numerous leukocytes |
| 41.1 | 3 | 3 | + | 2.5 | 2.5 | + | 2.5 | 2.5 | + | 2 | 2.5 | + | Brain: scattered ind neurons and axons; SC, SG: some neurons and ependyma; pancreas: several random acini (AEC) and ind AEC; spleen: numerous ind leukocytes |
| 41.2 | 2.5 | NE | NE | 2.5 | 3 | + | 3 | 3 | + | 2 | 2 | + | SC, SG: a few neurons; pancreas: numerous random pos acini (AEC); spleen: numerous ind leukocytes |
| 41.3 | 3 | 3 | + | 2 | 2 | + | 3 | 3 | + | 2 | 2.5 | + | Brain: a few ind neurons; SC: rare neuron (weak), ependymal cells; pancreas: numerous random pos acini (AEC); spleen: numerous ind leukocytes |
| 41.4 | 2.5 | 2.5 | + | 1.5 | 2 | + | 0.5 | 2.5 | + | 0.5 | 1.5 | + | Brain: a few random ind pos neurons; SC, SG: ind pos neurons, ependymal cells; pancreas: several random pos acini (AEC); spleen: several ind leukocytes |
| 41.5 | 3 | 3 | (+) | 3 | 3 | + | 3 | 3 | + | NE | NE | NE | Brain: rare ind pos neuron; SC, SG: a few pos neurons, ependymal cells; pancreas: several random pos acini (AEC) |

| | | | | | | | | | | | | | |
|---|---|---|---|---|---|---|---|---|---|---|---|---|---|
| 42 | 3 | 3 | + | 2 | 2 | + | 2.5 | 2.5 | (+) | 2 | 1.5 | (+) | Brain, SC: dot-like expression in periphery of ind neurons; pancreas: scattered ind AEC; spleen: scattered ind leukocytes |
| 43 | 2 | 2.5 | + | 2 | 2 | + | 2.5 | NE | NE | 2 | 2 | (+) | Brain, SC: ind pos neurons (cytoplasmic or peripheral reaction), ependymal cells; spleen: rare ind leukocytes |

Legend: Scoring was performed on tissue sections stained with HE or by IHC for viral NP expression, using a scale from 0 to 3 (0: no IB/reaction; 3: very abundant IB/widespread expression; with 0.5 step intermediate values). GP – determination of the detection of viral GP expression in sections stained by immunohistochemistry (- no reaction; (+) very limited reaction; + distinct reaction);

EC – epithelial cells; ggl – spinal ganglion; HE – IB score based on hematoxylin and eosin stained section; ind – individual; NE – not examined (or stained sections not suitable for evaluation due to extensive endogenous pigment deposition); NP – IB score based on sections stained by immunohistochemistry for the expression of viral nucleoprotein; SC – spinal cord; SG – spinal ganglia
